# The endonuclease Cue2 cleaves mRNAs at stalled ribosomes during No Go Decay

**DOI:** 10.1101/671099

**Authors:** Karole N. D’Orazio, Colin Chih-Chien Wu, Niladri Sinha, Raphael Loll-Krippleber, Grant W. Brown, Rachel Green

## Abstract

Translation of problematic sequences in mRNAs leads to ribosome collisions that trigger a sequence of quality control events including ribosome rescue, degradation of the stalled nascent polypeptide via the Ribosome-mediated Quality control Complex (RQC), and targeting of the mRNA for decay (No Go Decay or NGD). Previous studies provide strong evidence for the existence of an endonuclease involved in the process of NGD though the identity of the endonuclease and the extent to which it contributes to mRNA decay remain unknown. Using a reverse genetic screen in yeast, we identify Cue2 as the conserved endonuclease that is recruited to stalled ribosomes to promote NGD. Ribosome profiling and biochemistry provide strong evidence that Cue2 cleaves mRNA within the A site of the colliding ribosome. Finally, we show that NGD primarily proceeds via Xrn1-mediated exonucleolytic decay. Cue2-mediated endonucleolytic decay normally constitutes a secondary decay pathway, but becomes a major contributor in cells depleted of factors required for the resolution of stalled ribosome complexes (the RQT factors including Slh1). Together these results provide insights into how multiple decay processes converge to process problematic mRNAs in eukaryotic cells.

**One Sentence Summary:** Cue2 is the endonuclease that cleaves mRNA at ribosome stall sites.

## Introduction

Translation is a highly regulated process in which ribosomes must initiate, elongate, and terminate accurately and efficiently to maintain optimal protein levels. Disruptions in the open reading frames of mRNAs cause ribosome stalling events, which trigger downstream quality control pathways that carry out ribosome rescue, nascent peptide degradation (via the Ribosome-mediated Quality control Complex or RQC) and mRNA decay (No Go Decay or NGD) (Brandman & Hegde, 2016; Simms, Thomas, & Zaher, 2017). Recent work in eukaryotes has revealed that ribosome collisions on problematic mRNAs create a unique interface on the aligned 40S subunits that serves as a substrate for E3 ubiquitin ligases such as Hel2 and Not4, and the RQC-trigger (RQT) complex, comprised of factors Slh1, Cue2 and Rqt4; together these factors are thought to trigger downstream quality control (Ferrin & Subramaniam, 2017; Garzia et al., 2017; Ikeuchi et al., 2019; Juszkiewicz et al., 2018; Juszkiewicz & Hegde, 2017; Matsuo et al., 2017; Simms, Yan, & Zaher, 2017; Sundaramoorthy et al., 2017). A failure to process such colliding ribosomes and their associated proteotoxic nascent peptide products results in broad cellular distress, made evident by the strong conservation of these pathways (Balchin, Hayer-Hartl, & Hartl, 2016).

Previous genetic screens and biochemistry have identified key factors involved in the recognition of stalling or colliding ribosomes and in targeting the nascent polypeptide to the RQC (Brandman et al., 2012; Kuroha et al., 2010; D. P. Letzring, Wolf, Brule, & Grayhack, 2013). As these earliest screens focused on the identification of factors that stabilized reporter protein expression, factors involved in regulating mRNA decay by NGD remain largely unknown. The hallmark of NGD (Frischmeyer et al., 2002; van Hoof, Frischmeyer, Dietz, & Parker, 2002) is the presence of endonucleolytic cleavage events upstream of the ribosome stalling sequence as first detected by northern analysis (Doma & Parker, 2006) and more recently by high resolution ribosome profiling or sequencing approaches (Arribere & Fire, 2018; N. Guydosh & Green, 2017; N. R. Guydosh, Kimmig, Walter, & Green, 2017; Simms, Yan, et al., 2017). Importantly, these mRNA cut sites depend on ribosome collisions and the consequent polyubiquitination of the 40S subunit by the yeast protein Hel2 (Ikeuchi et al., 2019; Simms, Yan, et al., 2017). Multiple studies in the field collectively position NGD cleavage events within the vicinity of colliding ribosomes (Arribere & Fire, 2018; N. Guydosh & Green, 2017; N. R. Guydosh et al., 2017; Ibrahim, Maragkakis, Alexiou, & Mourelatos, 2018; Ikeuchi et al., 2019; Simms, Kim, Yan, Qiu, & Zaher, 2018; Simms, Yan, et al., 2017).

Interestingly, an endonuclease has been implicated in a related mRNA decay pathway, Nonsense Mediated Decay (NMD), in metazoans. During NMD, the ribosome translates to a premature stop codon (PTC), where an initial endonucleolytic cleavage event is carried out by a critical PIN-domain containing endonuclease, SMG6 (Glavan, Behm-Ansmant, Izaurralde, & Conti, 2006); sequencing experiments suggest that SMG6 cleavage occurs in the A site of PTC-stalled ribosomes (Arribere & Fire, 2018; Lykke-Andersen et al., 2014). In *C. elegans*, there is evidence that the initial SMG6-mediated endonucleolytic cleavage leads to iterated cleavages upstream of the PTC similar to those characterized for the NGD pathway in yeast (Arribere & Fire, 2018). The idea that the various decay pathways may act synergistically is intriguing. Homologs of SMG6 (NMD4/EBS1 in yeast) (Dehecq et al., 2018) and other PIN domain-containing proteins, as well as other endonuclease folds, exist throughout eukarya and anecdotally have been evaluated by the field as potential candidates for functioning in NGD, though there are no reports to indicate that any function in this capacity. The identity of the endonuclease responsible for cleavage in NGD remains unknown.

The presumed utility of endonucleolytic cleavage of a problematic mRNA is to provide access to the mRNA for the canonical exonucleolytic decay machinery that broadly regulates mRNA levels in the cell. In yeast, Xrn1 is the canonical 5’ to 3’ exonuclease which, after decapping of the mRNA, degrades mRNA from the 5’ end; importantly, Xrn1 normally functions co-translationally such that signals from elongating ribosomes might be relevant to its recruitment (W. Hu, Sweet, Chamnongpol, Baker, & Coller, 2009; Pelechano, Wei, & Steinmetz, 2015). Additionally, the exosome is the 3’ to 5’ exonuclease which,after deadenylation, degrades mRNAs from the 3’ end and is recruited by the SKI auxiliary complex consisting of Ski2/Ski3/Ski8 and Ski7 (Halbach, Reichelt, Rode, & Conti, 2013). While Xrn1-mediated degradation is thought to be the dominant pathway for most general decay in yeast (Anderson & Parker, 1998), the exosome has been implicated as critical for many degradation events in the cell including those targeting prematurely polyadenylated mRNAs (these mRNAs are usually referred to as Non-Stop Decay (NSD) targets) (Frischmeyer et al., 2002; Tsuboi et al., 2012; van Hoof et al., 2002). In metazoans, it is less clear what the relative contributions of Xrn1 and the exosome are to the degradation of normal cellular mRNAs. How the endonucleolytic and canonical exonucleolytic decay pathways coordinate their actions on problematic mRNAs remains unknown.

Here we present a reverse genetic screen in *S. cerevisiae* that identifies Cue2 as the primary endonuclease in NGD. Using ribosome profiling and biochemical assays, we show that Cue2 cleaves mRNAs in the A site of collided ribosomes, and that ribosomes which accumulate at these cleaved sites are rescued by the known ribosome rescue factor Dom34 (N. R. Guydosh & Green, 2014; Shoemaker, Eyler, & Green, 2010). We further show that stall-dependent endonucleolytic cleavage represents a relatively minor pathway contributing to the decay of the problematic mRNA reporter, while exonucleolytic processing by the canonical decay machinery, in particular Xrn1, plays the primary role. The Cue2-mediated endonucleolytic cleavage activity is substantially increased in genetic backgrounds lacking the factor Slh1, a known component of the RQT complex (Matsuo et al., 2017), suggesting that the relative contribution of this pathway could increase in different environmental conditions. Our final model provides key insights into what happens in cells upon recognition of stalled ribosomes on problematic mRNAs, and reconciles how both endo- and exonucleolytic decay act synergistically to resolve these dead-end translation intermediates.

## Results

### Screening for factors involved in NGD

To identify factors that impact the degradation of mRNAs targeted by NGD, we developed a construct that directly reports on mRNA levels. Previous genetic screens in yeast (Brandman et al., 2012; Kuroha et al., 2010; D. P. Letzring et al., 2013) were based on reporters containing a stalling motif in an open reading frame (ORF). As a result, these screens primarily revealed machinery involved in recognition of stalled ribosomes and in degradation of the nascent polypeptide, but missed factors involved in mRNA decay. In our reporters (*GFP-2A-FLAG-HIS3*), the protein output for the screen (GFP) is decoupled from the stalling motif positioned within *HIS3* by a 2A self-cleaving peptide sequence (Di Santo, Aboulhouda, & Weinberg, 2016; Sharma et al., 2012) (Fig. 1A). Because GFP is released before the ribosome encounters the stalling sequence within the *HIS3* ORF, its abundance directly reflects the reporter mRNA levels and translation efficiency independent of the downstream consequences of nascent peptide degradation. These reporters utilize a bidirectional galactose inducible promoter such that an *RFP* transcript is produced from the opposite strand and functions as an internal-control for measurement of general protein synthesis.

**Fig. 1:**
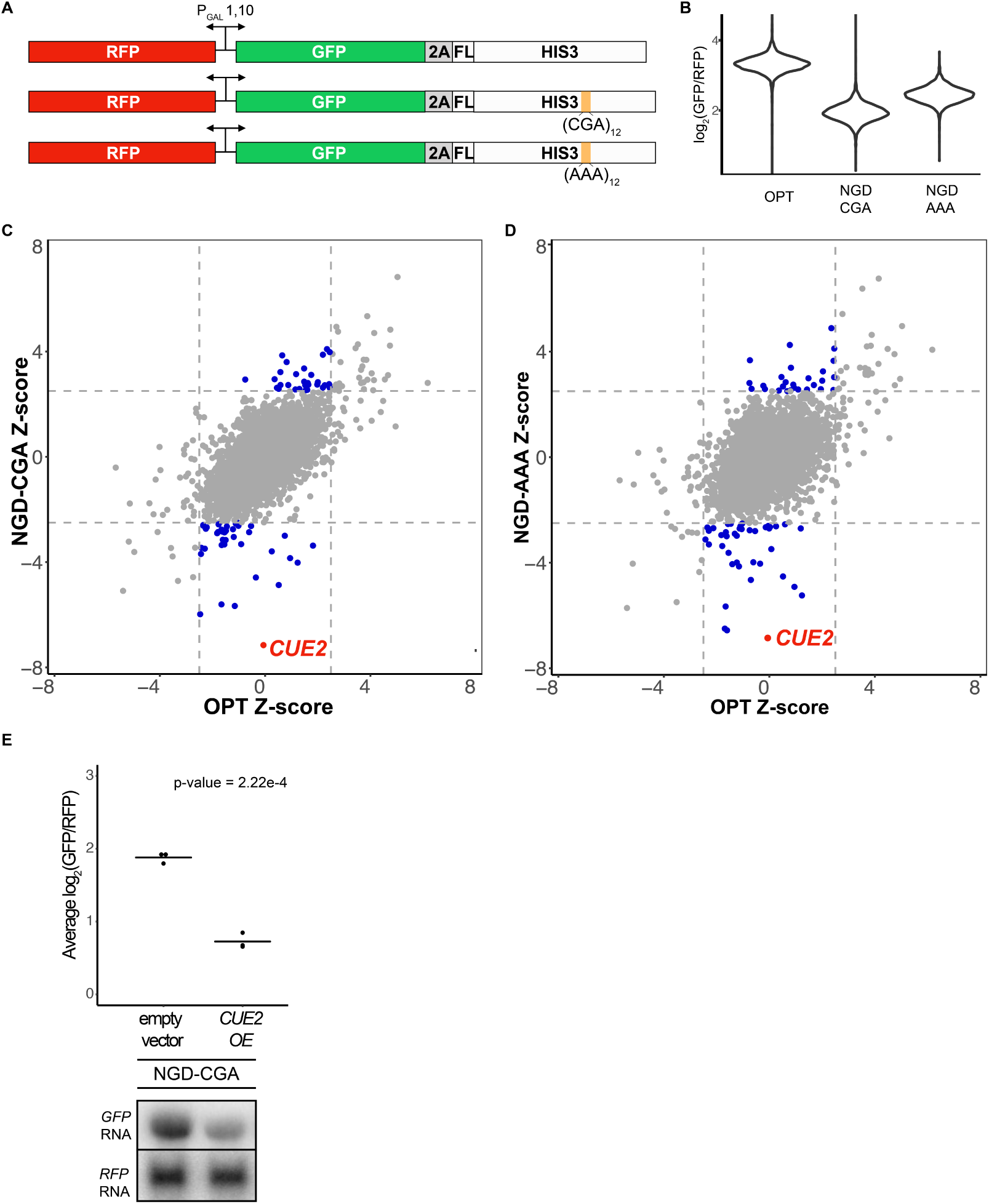
Yeast overexpression screens identify a novel factor involved in NGD. **(A)** Schematic of reporters used in genetic screens. **(B)** Normalized reporter GFP levels. Violin plots show flow cytometry data from > 4000 WT cells containing the indicated reporter. **(C-D)** Plots of Z-scores overexpression screens comparing the NGD-CGA and NGD-AAA reporters to OPT. Z-scores reflecting the significance of log_2_(GFP/RFP) values from each strain are plotted against each other for the two reporters. Dashed lines represent cutoffs at a Z-score greater than 2.5 or less than −2.5 for each reporter. Blue dots represent deletion strains that have a Z-score value outside the cut-off for the NGD reporters, but not the OPT. Red dots represent strains of interest in this study. **(E)** Validation data for overexpression screen hit *CUE2*. Top, means for 3 individual flow cytometry experiments are plotted for the NGD-CGA reporter strain without *CUE2* overexpression (left), and with *CUE2* overexpression (right). Bottom, northern blots of steady state mRNA levels for the same strains.

The screen utilized two different NGD constructs with stalling motifs inserted into the *HIS3* gene: the first contains 12 CGA codons (NGD-CGA) which are decoded slowly (Kuroha et al., 2010; Daniel P. Letzring, Dean, & Grayhack, 2010) by the low-copy ICG-tRNA^Arg^ and the second contains 12 AAA codons (NGD-AAA) that mimic polyA tail and are known to trigger ribosome stalling and mRNA quality control in both yeast and mammalian systems (Arthur et al., 2015; Frischmeyer et al., 2002; Garzia et al., 2017; N. Guydosh & Green, 2017; Ito-Harashima, Kuroha, Tatematsu, & Inada, 2007; Juszkiewicz & Hegde, 2017; Sundaramoorthy et al., 2017; van Hoof et al., 2002) (Fig. 1A). The (CGA)_12_ and (AAA)_12_ inserts result in robust 3-fold and 2-fold decreases in the GFP/RFP ratio, respectively, compared to the no insert (optimal, OPT) control (Fig. 1B). Similar changes are also seen in GFP levels by western blot and in full-length mRNA levels by northern blot (Fig. S1A). As previously reported, deletion of the exosome auxiliary factor gene *SKI2* stabilized the 5’ decay fragment resulting from endonucleolytic cleavage associated with NGD reporters (Fig. S1A) (Doma & Parker, 2006). Finally, stalling during the synthesis of the FLAG-His3 fusion protein in the two stalling reporters (NGD-CGA and NGD-AAA) leads to degradation of the nascent peptide (Fig. S1A, FLAG panel).

We used high-throughput reverse genetic screens and reporter-synthetic genetic array (R-SGA) methods (Fillingham et al., 2009; Tong et al., 2001) to evaluate the effects of overexpressing annotated genes on GFP expression. We began by crossing strains carrying the control (OPT) and the two different no-go decay reporters (NGD-CGA and NGD-AAA) described in Fig. 1A into the *S. cerevisiae* overexpression library (Douglas et al., 2012; Giaever et al., 2002; Y. Hu et al., 2007). For each overexpression screen, we isolated diploid strains containing both the overexpression plasmid and our reporter. Selected strains were transferred to galactose-rich plates and the GFP and RFP levels were evaluated by fluorimetry. We plotted the results from the screens individually, comparing Z-scores for the log_2_(GFP/RFP) signals from each NGD reporter strain to Z-scores from the corresponding strain carrying the OPT reporter (Fig. 1C-1D). Normalization with RFP intensity was used to eliminate non-specific factors that impact expression of both RFP and GFP.

The overexpression screen revealed a set of candidate genes that modulate GFP levels for the NGD reporters relative to the OPT reporter (Fig. 1C-1D and Table S2). Broadly, we see a stronger overlap in candidate overexpression genes among the NGD reporters than for either NGD reporter compared to the OPT reporter (Fig. S1B). By far, the strongest outlier by Z-score causing reduced GFP expression for both NGD reporters, without affecting the OPT reporter, resulted from overexpression of the gene *CUE2* (Fig. 1C-1D). Flow cytometry and northern analysis confirm that NGD-CGA reporter mRNA levels are substantially reduced upon *CUE2* overexpression whereas the control RFP transcript is not affected (Fig. 1E and S1C).

### *CUE2* domain structure and homology modeling

The domain structure of Cue2 reveals it to be a promising candidate for the missing endonuclease for NGD. *CUE2* contains two CUE (coupling of ubiquitin to ER degradation) ubiquitin-binding domains at the N-terminus (Kang et al., 2003) followed by a structurally undefined linker region and an SMR (small MutS-related) hydrolase domain at the C-terminus (Fig. 2A). We performed alignments of the CUE and SMR domains of Cue2 using structure-based alignment tools (Fig. S2A and S2B, respectively) and found putative homologs in various kingdoms of life (including the human NEDD4-binding protein 2, N4BP2). While the SMR domain of MutS family enzymes canonically function as DNA-nicking hydrolases (Fukui & Kuramitsu, 2011), the SMR domain of SOT1 in plants exhibits RNA endonuclease activity (Zhou et al., 2017). Additionally, SMR domains show structural similarity to bacterial RNase E (Fukui & Kuramitsu, 2011). Alignments of the SMR domains of these and other proteins enabled us to identify conserved residues (Fig. 2B), some of which are known to be critical for RNA endonuclease activity in the plant enzyme (Zhou et al., 2017).

**Fig. 2:**
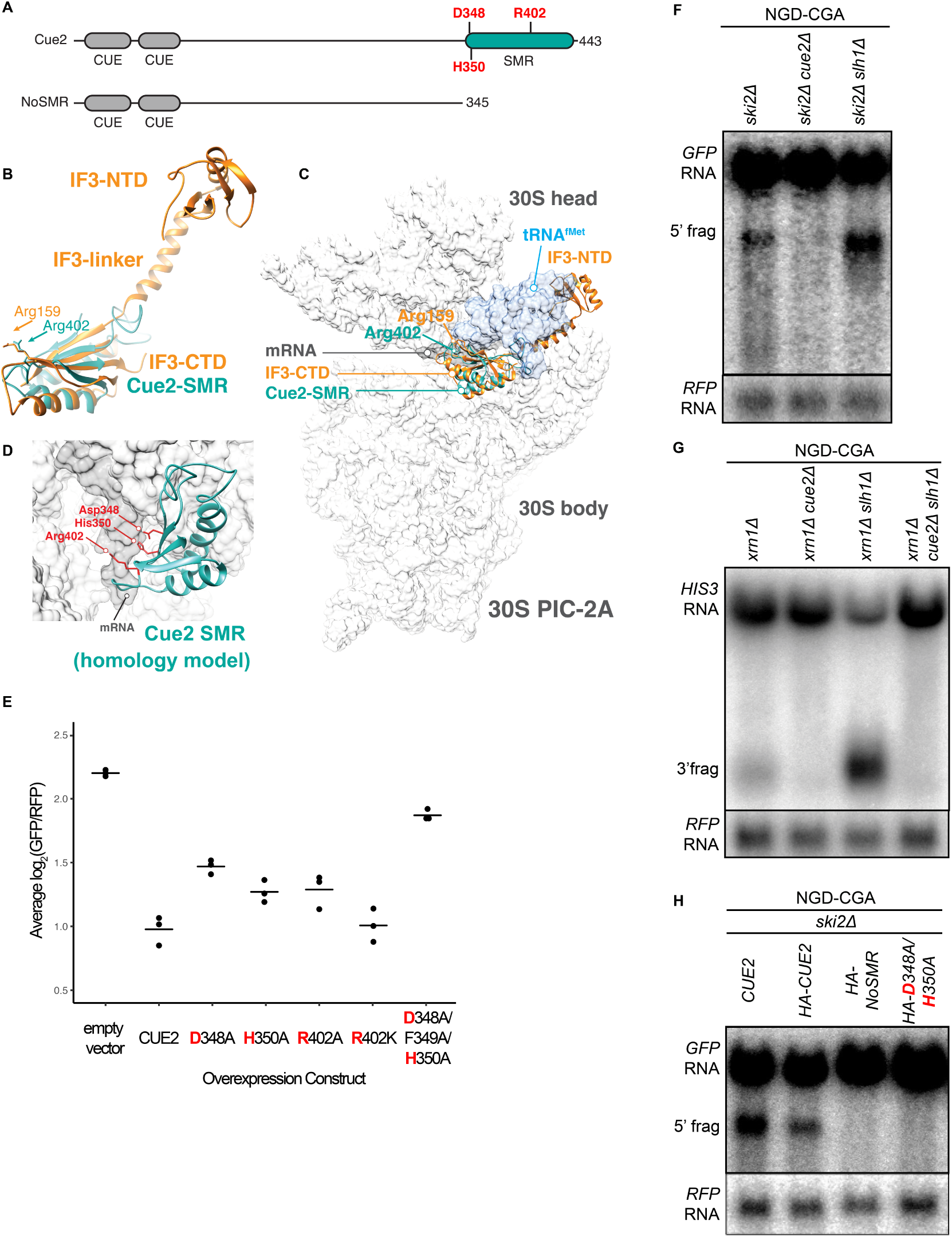
Cue2 is the endonuclease what cleaves mRNA during No Go Decay. **(A)** Domain organization of *S. cerevisiae* Cue2, with the “NoSMR” mutant schematic below the WT *CUE2*. **(B)** Superimposition of homology model of Cue2 SMR domain (cyan) (templated on human N4BP2 SMR; PDB: 2VKC (Diercks et al., 2008)) on full-length IF3 (orange, PDB: 5LMQ (Hussain et al., 2016)). Positions of Arg-159 of IF3-CTD and Arg-402 of Cue2-SMR are indicated. **(C)** Putative positioning of Cue2-SMR in the context of IF3 and tRNA^fMet^ bound to the small ribosome subunit. 30S PIC-2A (as described in Hussain et al., 2016 (Hussain et al., 2016)) is light gray; mRNA is dark gray; tRNA^fMet^ is blue; IF3 is orange; Cue2-SMR is cyan. **(D)** Cue2-SMR homology model in the same orientation as seen in Fig. S2E, showing putative positions of Asp-348, His-350 and Arg-402 with 30S pre-initiation complex (PIC) with tRNA^fMet^ density subtracted (PDB: 5LMQ, State 2A (Hussain et al., 2016)). **(E)** Mutation analysis of Cue2 SMR domain. Log_2_(GFP/RFP) flow data with the indicated overexpression constructs are shown. P-values for the utant *HA-CUE2* overexpression genes as compared to the wildtype *HA-CUE2* overexpression gene are 0.007195, 0.02602, 0.03857, 0.7788, 0.002127 and 0.002869, respectively. **(F-G)** Northern blot analysis of the indicated strains, with full-length mRNA and the 5’ **(F)** and 3’ **(G)** fragments labeled; RFP probed for normalization. **(H)** Northern blot analysis of NGD-CGA cleavage fragments in the *ski2Δ* strain with the indicated changes to the endogenous *CUE2* locus; RFP probed for normalization.

Heuristic searches of the structurally defined SMR domain of the mammalian homolog of Cue2 (N4BP2) (Diercks et al., 2008) against known structures in the Protein Data Bank found it to be structurally homologous to the C-terminal domain (CTD) of bacterial Initiation Factor 3 (Biou, Shu, & Ramakrishnan, 1995) (Fig. 2B and Fig. S2C-S2D). During initiation in bacteria, IF3 binds to the 30S pre-initiation complex (PIC) and helps position the initiator tRNA at the AUG start codon; the CTD of IF3 binds in close proximity to the P and A sites of the small subunit, closely approaching the mRNA channel (Hussain, Llácer, Wimberly, Kieft, & Ramakrishnan, 2016). We aligned the Cue2-SMR homology model with the structure of the IF3-CTD on the ribosome and observed that conserved residues D348, H350, and R402 are positioned along the mRNA channel in this model (Fig. 2C-2D and Fig. S2E-S2F) and thus represent potential residues critical for endonucleolytic cleavage activity; additionally, R402 had previously been implicated in the RNA cleavage activity of other SMR domains (Zhou et al., 2017).

### Characterizing roles of *CUE2* in NGD *in vivo*

We performed several different experiments to ask if these conserved residues are necessary for Cue2 function. First, we generated mutations in HA-tagged Cue2 and performed flow cytometry on the NGD-CGA reporter under overexpression conditions for the different Cue2 variants. We find that individually mutating the conserved residues D348, H350, and R402 to alanine (A) causes a modest increase in GFP expression relative to *CUE2* WT, whereas the R402K mutation (which maintains a positively charged amino acid) has no discernible effect (Fig. 2E). Mutating residues 348 through 350, which includes the conserved residues D348 and H350, nearly restores GFP reporter signal in this assay (Fig. 2E). Importantly, each of these variants is expressed at similar levels (Fig. S2G). These data suggest a potential role for residues D348, H350, and R402 in the SMR domain of Cue2 in reducing levels of problematic mRNAs.

We next asked if Cue2 is necessary for the previously documented endonucleolytic cleavage of the NGD reporter transcripts. In order to visualize the mRNA fragments resulting from cleavage, we deleted either the 3’ - 5’ mRNA decay auxiliary factor, *SKI2*, or the major 5’ - 3’ exonuclease, *XRN1*, to stabilize the 5’- and 3’-fragments, respectively (Doma & Parker, 2006) (Fig. 2F; lane 1 and 2G; lane 1). Upon deletion of *CUE2* in the appropriate yeast background, we see that both of these decay intermediates disappear (Fig. 2F; lane 2 and 2G; lane 2).

In order to further investigate the role of the specific amino acids proposed to be catalytically critical, we modified *CUE2* at its endogenous chromosomal locus with an HA tag, to allow us to follow native *CUE2* protein levels, and generated mutations at the endogenous *CUE2* locus in a *ski2Δ* background. As a first test, we deleted the C-terminal portion of *CUE2* comprising the SMR domain and revealed a complete loss of endonucleolytic cleavage (Fig 2H, lane 3). To further refine this analysis, we made combined mutations in *CUE2* at D348 and H350 and observed a complete abolishment of endonucleolytic cleavage activity. Western analysis confirmed that protein expression levels from the CUE2 locus are equivalent for these different protein variants (Figure S2H). These data provide strong support for a hydrolytic role for Cue2 in the endonucleolytic cleavage of problematic mRNAs in the NGD pathway.

### Contribution of Cue2 to NGD is increased in specific genetic backgrounds

Surprisingly, deletion of *CUE2* in the WT, *ski2Δ* or *xrn1Δ* background does not demonstrably impact GFP or steady-state full-length mRNA levels (Fig. 3A-3B, Fig. 2F, or Fig. 2G, respectively) suggesting that the endonucleolytic decay pathway is not responsible for the majority of the three-fold loss in mRNA levels observed for this NGD reporter (relative to OPT). However, we see that deleting *XRN1* alone greatly restores NGD reporter mRNA levels while deleting *SKI2* alone has very little effect (Fig. 3A-3B). These results together suggest that the majority of mRNA degradation observed for these NGD reporters is mediated by canonical decay pathways, and mostly Xrn1.

**Fig. 3:**
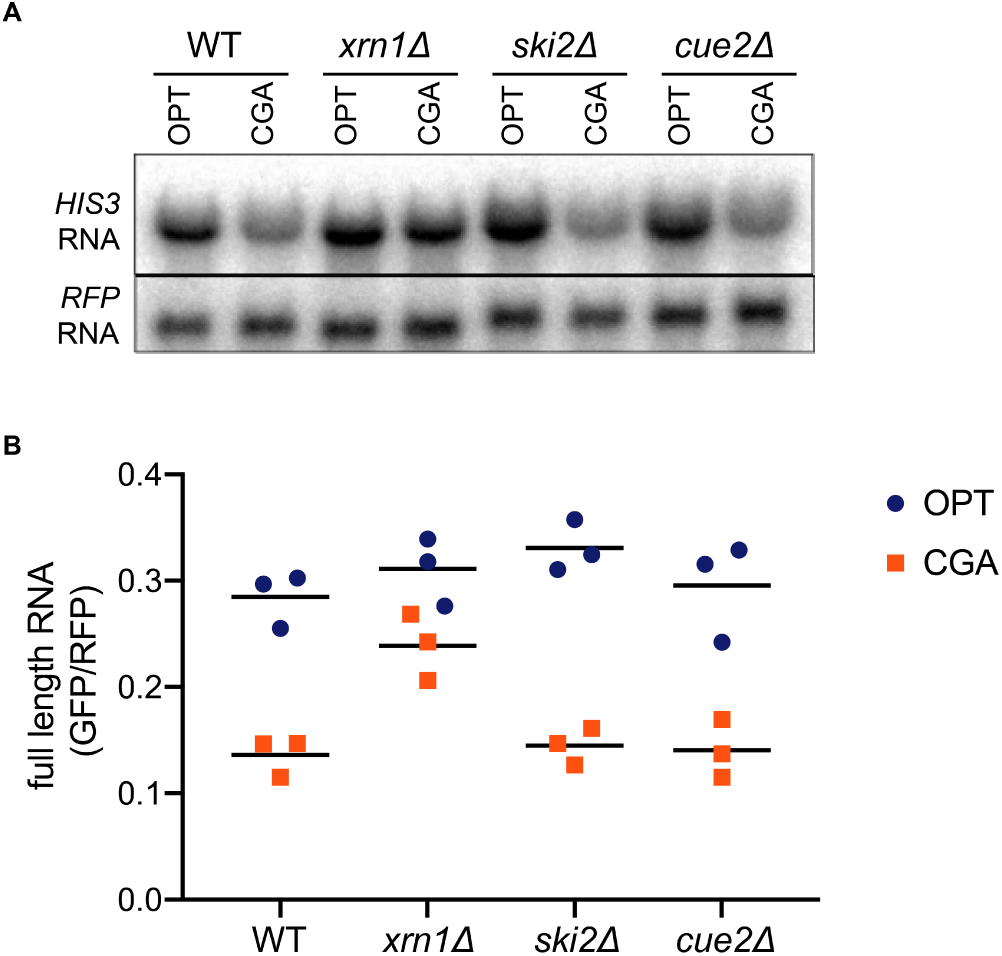
Canonical decay by Xrn1 is the major contributor to No Go Decay. **(A)** Northern blot analysis of full length OPT and NGD-CGA reporter levels in the indicated strains. **(B)** Quantitated northern blot signal from full length GFP/RFP mRNA levels for the indicated strain, in triplicate.

A key insight into the regulation of decay by these distinct mRNA decay pathways (endo- and exo-nucleolytic) came from examining decay intermediates in strains lacking the factor Slh1. Slh1 is a member of the Ribosome Quality Trigger complex that was previously implicated in regulating NGD (Ikeuchi et al., 2019; Matsuo et al., 2017). This large protein (∼200 kDa) is homologous to helicases implicated in RNA decay and splicing including Ski2 and Brr2, respectively (Johnson & Jackson, 2013). Slh1 contains two DEAD box motifs and mutations in them are known to disrupt targeting of the nascent peptide on problematic mRNAs to proteolytic decay by the RQC. Based on these observations, Slh1 has been proposed to directly or indirectly dissociate stalled ribosomes and to thereby target them for the RQC (Matsuo et al., 2017).

We found that in the *ski2Δ slh1Δ* and *xrn1Δ slh1Δ* strains, we see a sizeable build-up of endonucleolytically cleaved mRNA fragments (Fig. 2F; lane 3 and 2G; lane 3). Consistent with the recent study by Ikeuchi et al. (Ikeuchi et al., 2019), we also see smearing of the signal representative of shorter 5’ and longer 3’ fragments due to additional cleavage events occurring upstream of the primary stall site in the absence of *SLH1*. Importantly, we also observe a robust decrease in full-length mRNA levels upon deletion of *SLH1* (Fig. 2F; lane 3 and 2G; lane 3). These data suggest a model wherein Slh1 activity somehow competes with Cue2-mediated endonucleolytic cleavage. To ask whether the increased endonucleolytic cleavage seen in the *SLH1* deletion background is the result of Cue2 activity, we deleted *CUE2* in the *xrn1Δ slh1Δ* background and observed a complete loss of the decay intermediate and a restoration of full-length mRNA levels (Fig. 2G; lane 4). These data provide strong evidence that the action of Slh1 negatively regulates the assembly of a potent Cue2 substrate though the molecular mechanism for this regulation is not defined.

### Ribosome profiling provides high-resolution view of Cue2 cleavage sites

We next utilized ribosome profiling to assess the role and specificity of Cue2 in endonucleolytic cleavage of problematic mRNAs. Ribosome profiling was performed in various genetic backgrounds where the NGD-CGA reporter construct was included to follow the process of NGD on a well-defined mRNA substrate. The approach benefits from our ability to distinguish three distinct sizes of ribosome-protected mRNA fragment (RPF) that correspond to three different states of the ribosome during translation. 16 nucleotide (nt) RPFs correspond to ribosomes that have translated to the truncated end of an mRNA in the cell, usually an mRNA that has been endo- or exonucleolytically processed; these 16 nt RFPs are enriched in a *dom34Δ* strain since these are established targets for Dom34-mediated ribosome rescue (Arribere & Fire, 2018; N. Guydosh & Green, 2017; N. R. Guydosh & Green, 2014; N. R. Guydosh et al., 2017). 21 and 28 nt RPFs are derived from ribosomes on intact mRNA, with the two sizes corresponding to different conformational states of ribosomes in the elongation cycle (Lareau, Hite, Hogan, & Brown, 2014; Wu, Zinshteyn, Wehner, & Green, 2019). In our modified ribosome profiling library preparation (Wu et al., 2019), 21 nt RPFs represent classical/decoding ribosomes waiting for the next aminoacyl-tRNA whereas the 28 nt RPFs primarily represent ribosomes in a rotated/pre-translocation state. Throughout this study, we performed ribosome profiling in a *ski2Δ* background to prevent 3’-5’ exonucleolytic degradation by the exosome and enable detection of mRNA decay intermediates. We note that deleting *SKI2* generally stabilizes prematurely polyadenylated and truncated mRNAs in the cell, as seen previously (Tsuboi et al., 2012; van Hoof et al., 2002), therefore use of this background likely enhanced Cue2 cleavage.

In a first set of experiments, we compare the distributions of RPFs in *ski2Δ* strains to strains additionally lacking *DOM34*, or *DOM34* and *CUE2*; in each case, we separately consider the different RPF sizes (16, 21 and 28) to determine the state of the ribosome on these sequences. In *ski2Δ*, we observe a clear accumulation of ribosome density at the (CGA)_12_ stall site (in the 21 nt RPF track) in our reporter (Fig. 4A) consistent with ribosome stalling at this site. Looking first within the actual (CGA)_12_ codon tract, we see a quadruplet of 21 nt RPF peaks from the 2^nd^ to the 5^th^ CGA codon (middle panel). Because of difficulties in mapping reads to repetitive sequences such as (CGA)_12_, we also isolated RPFs from a minor (< 5% of total reads on reporter) population of stacked disome species (N. R. Guydosh & Green, 2014) and show that although some ribosomes move into the subsequent CGA codons, the principal ribosome accumulation occurs between the 2^nd^ to the 5^th^ codon (Fig. S3). While 21 nt RFPs are enriched on CGA codons, 28 nt RPFs are not (Fig. 4A, compare middle and bottom panels), indicating that most ribosomes found at the CGA codons are waiting to decode the next aminoacyl-tRNA; these observations are consistent with the fact that CGA codons are poorly decoded (Daniel P. Letzring et al., 2010). The 21 nt RPF quadruplet at the CGA cluster represents the stalled “lead” ribosome in its classical (unrotated) conformation, which is consistent with recent cryoEM structures of collided ribosomes (Ikeuchi et al., 2019; Juszkiewicz et al., 2018). About 30 nts upstream of these lead ribosomes, we observe the accumulation of 28 nt RPFs that correspond to colliding ribosomes in a rotated state (Fig. 4A, bottom panel), in agreement with the ribosome states recently reported in cryo-EM structures of collided disomes. In the *dom34Δ ski2Δ* strain (where 16 nt RPFs are stabilized), we observe a strong quadruplet of 16 nt RPF peaks exactly 30 nts upstream of the quadruplet of 21 nt RPFs (Fig. 4B), indicative of endonucleolytic cleavage events occurring upstream of the (CGA)_12_–stalled ribosomes. The precise 30 nt distance between the 16 nt RPFs and the downstream 21 nt RPFs is consistent with the measured distance between the A sites of two colliding ribosomes.

**Fig. 4:**
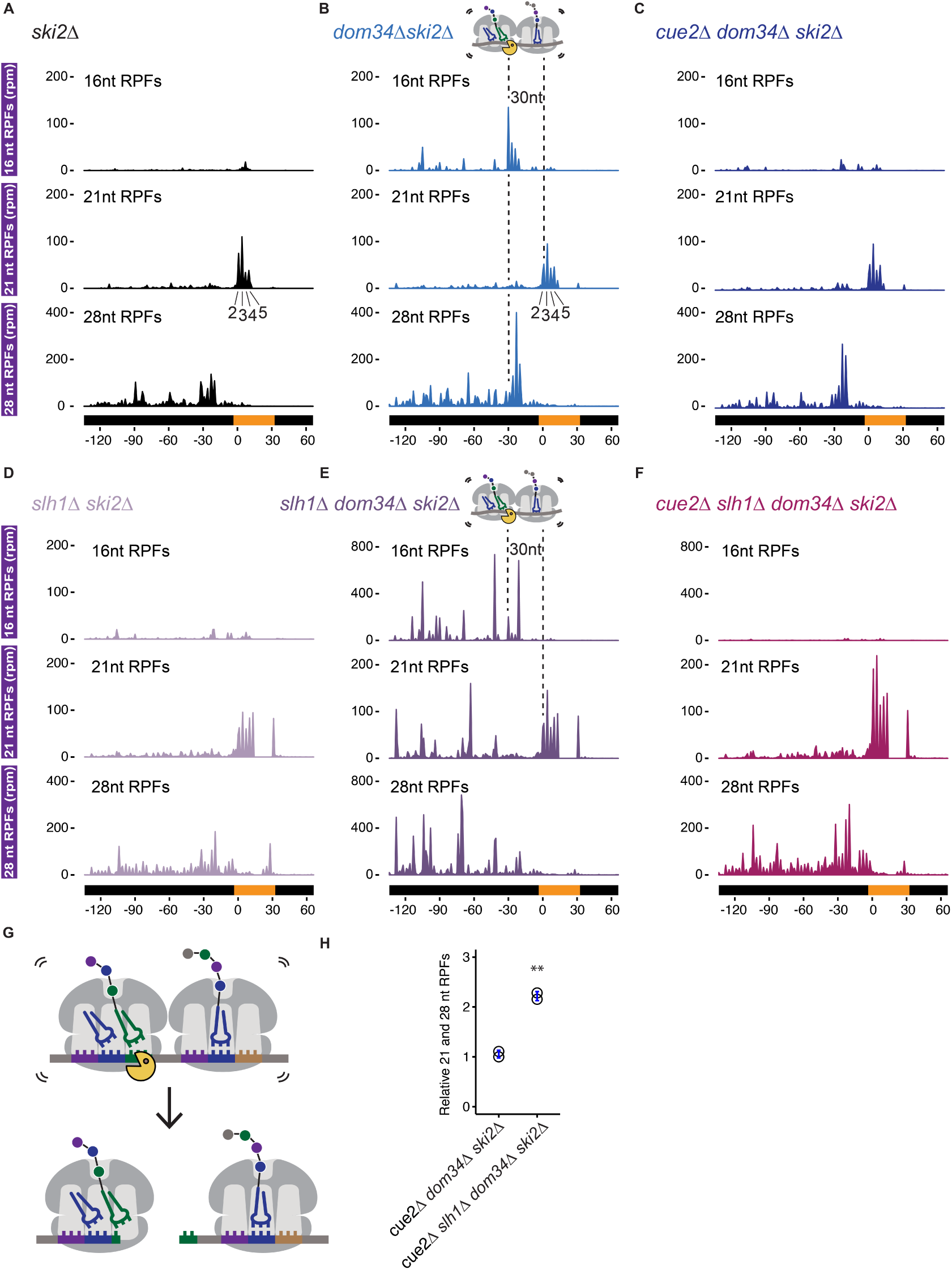
Ribosome profiling analysis of NGD on reporter mRNAs in various genetic backgrounds. **(A-F)** 16, 21 and 28 nt RPFs mapped to NGD-CGA reporter in *ski2Δ* **(A)**, *slh1Δ ski2Δ* **(B)**, *dom34Δ ski2Δ* **(C)**, *cue2Δ dom34Δ ski2Δ* **(D)**, *slh1Δ dom34Δ ski2Δ* **(E)**, and *cue2Δ slh1Δ dom34Δ ski2Δ* **(F)** strains with schematic depicting Cue2-mediated cleavage in the A sites of collided ribosomes. **(G)** Schematic of the precise Cue2 cleavage location on the mRNA, relative to collided ribosomes. **(H)** Comparison of the combined 21 and 28 nt ribosome occupancies on GFP, from 300 nt upstream of the (CGA)_12_ to the end of the (CGA)_12_ sequence, normalized to RFP for the indicated strains (n=2). p-value from Student’s t-test is indicated by asterisks. **, p< 0.01.

To address the role of Cue2 as the endonuclease, we examined the same collection of RPFs in the *cue2Δ dom34Δ ski2Δ* strain. In this background, the 16 nt RPFs indicative of endonucleolytic cleavage are dramatically reduced in abundance while the distributions of 21 and 28 nt RPFs are largely unaffected (Fig. 4C). These data argue that endonucleolytic cleavage occurs precisely in the A site of the collided, rotated ribosome and requires Cue2. A high-resolution view of the site of cleavage within the A site codon is found in the model in Figure 4G. These observations are broadly consistent with recent RACE analyses (Ikeuchi et al., 2019) and with our structural homology modeling placing Cue2 in the decoding center of the ribosome (Fig. 2C and 2D).

Based on our observations that deletion of Slh1 increases utilization of Cue2 for processing of problematic mRNAs, we next examined the distribution of RPFs on the NGD-CGA reporter in yeast strains lacking *SLH1*. In the *Δslh1 ski2Δ* strain, the pattern of RPF distributions is broadly similar to what we observed in the *ski2Δ* strain (Fig. 4D compared to 4A): 21 nt RPFs accumulate on the 2^nd^ through 5^th^ CGA codons and 28 nt RPFs accumulate 30 nts behind the lead ribosome; we also see modest accumulation of RPFs downstream of the stalling sequence as previously reported (Fig. 4D) (Sitron, Park, & Brandman, 2017). *SLH1* deletion does not stabilize 16 nt RPFs, and therefore, likely does not act on cleaved Cue2-substrates. When we delete *DOM34* in the *slh1Δ ski2Δ* background (*slh1Δ dom34Δ ski2Δ*), we see a dramatic accumulation of 16 nt RPF peaks (again, as a quadruplet, though with an altered relative distribution) behind the lead ribosomes (Fig. 4E compared to 4B, top panels, note the larger y-axis scale in Fig. 4E and 4F compared to 4B and 4C). These data are consistent with the substantial increase in cleavage we see via northern blot (Fig. 2F and 2G) and previous findings that deletion of the RQT complex causes somewhat distinct cleavage events (Ikeuchi et al., 2019). Importantly, deletion of *CUE2* in the *slh1Δ dom34Δ ski2Δ* background leads to a near complete loss of the 16 nt RPFs (Fig. 4F, top panel) and an enrichment of 21 and 28 nt RPFs.

Lastly, consistent with earlier studies showing that endonucleolytic cleavage events associated with NGD depend on the E3 ligase Hel2 (Ikeuchi et al., 2019), we show that deletion of *HEL2* in both *dom34Δ ski2Δ* and *slh1Δ dom34Δ ski2Δ* backgrounds leads to a complete loss of 16 nt RPFs (Fig. S4A and S4B, respectively).

### *In vitro* reconstitution of Cue2 cleavage on isolated colliding ribosomes

To test if Cue2 cleaves mRNA directly, we performed *in vitro* cleavage assays with the heterologously expressed SMR domain (and R402A mutant) of Cue2 and purified colliding ribosomes. To isolate the colliding ribosomes, cell-wide ribosome collision events were induced using low dose cycloheximide treatment of growing yeast (Simms, Yan, et al., 2017). We both optimized the yield and ensured that the purified Cue2-SMR domain was the only source of Cue2 in our experiment by isolating ribosomes from the *slh1Δ dom34Δ cue2Δ ski2Δ* strain; these cells also carried and expressed the NGD-CGA reporter. Since collided ribosomes are resistant to general nucleases (Juszkiewicz et al., 2018), MNase digestion of lysates collapses elongating ribosomes from polysomes to monosomes but spares those with closely packed (nuclease resistant) ribosomes (Fig. S5). As anticipated, only cells treated with low doses of cycloheximide yielded a substantial population of nuclease resistant, or collided, ribosomes after MNase digestion (Fig. S5, black trace). We isolated nuclease resistant trisomes from a sucrose gradient to serve as the substrate for our *in vitro* reconstituted cleavage experiment.

We purified the isolated SMR domain of Cue2 based on our *in vivo* evidence that this domain represents the functional endonuclease portion of the protein. We added the purified SMR domain of Cue2 to the isolated nuclease resistant trisomes and resolved the products of the reaction on a sucrose gradient that optimally resolves trisomes from monosomes and disomes. While we initially isolated trisomes, it is clear that the ‘untreated’ sample (black trace) contains trisomes, as well as disomes and monosomes; this may be the result of cross-contamination of those peaks during purification or the instability of the trisome complex. Nevertheless, on addition of Cue2-SMR (WT) (pink trace), we observe a substantial loss of trisomes and a corresponding increase in monosomes and disomes (Figure 5A), while on addition of the Cue2 SMR-R402A mutant (orange trace), we observe a more modest decrease in trisomes (Figure 5A). These results provide initial *in vitro* evidence that the SMR (hydrolase) domain of Cue2 cleaves the mRNA within a stack of colliding ribosomes.

**Fig. 5:**
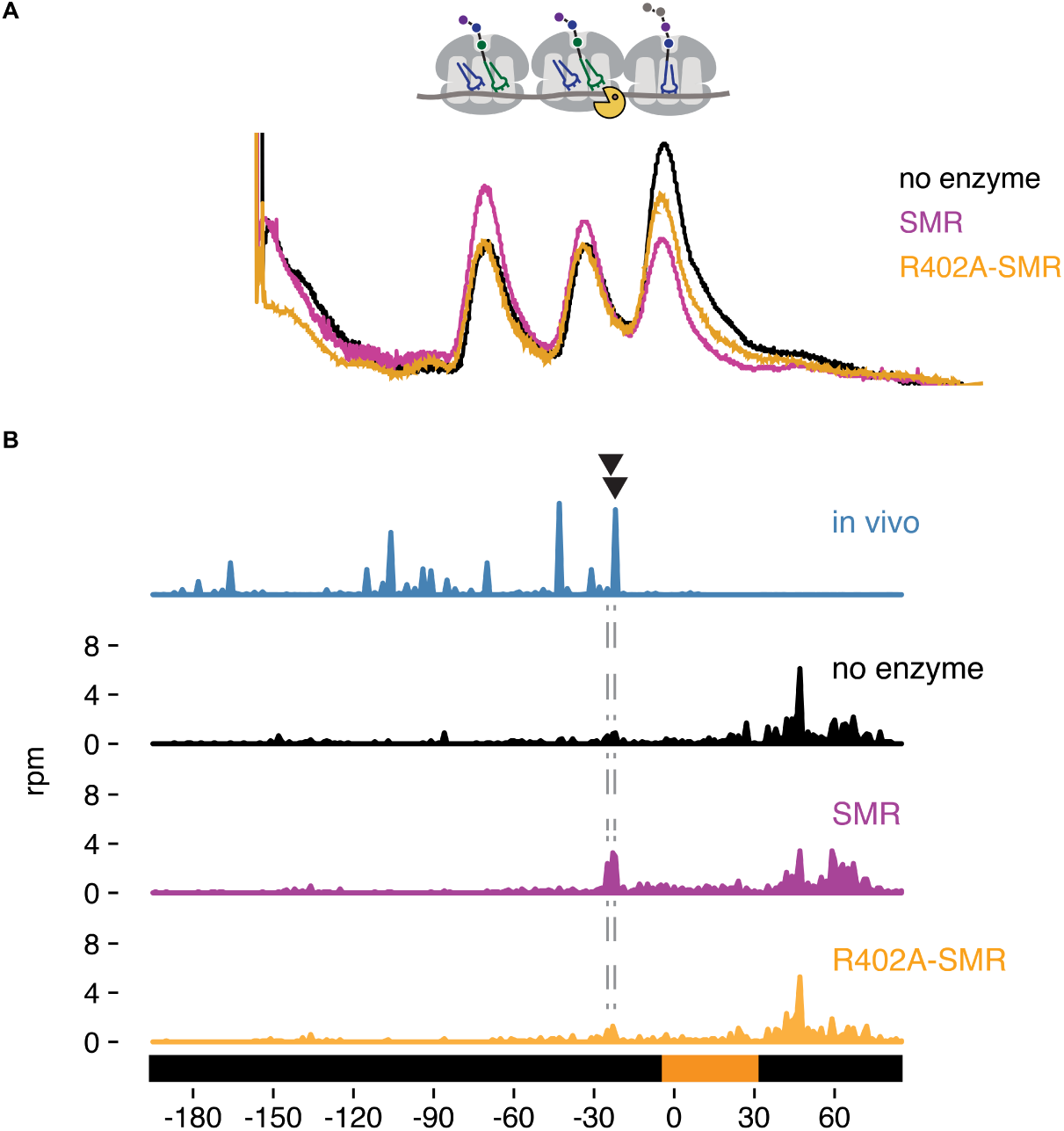
*In vitro* cleavage of purified Cue2-SMR. **(A)** Sucrose gradient of nuclease resistant trisomes, treated with no enzyme (black), the SMR domain of Cue2 (pink), or the SMR-R402A mutant domain (orange). **(B)** *In vivo* 16 nt RPFs from the *slh1Δ dom34Δ ski2Δ* strain (cyan) compared to the 60-65 nt RPFs from the combined fractions (mono-, di-and trisomes) treated with no enzyme (black), SMR domain (pink), or R402-SMR (orange) *in vitro*. Arrowheads indicate the positions where the *in vitro* coincide with the *in vivo* cleavages.

To extend these observations, we mapped more precisely the *in vitro* cleavage sites of the SMR domain by sequencing the ribosome-protected mRNA fragments from the untreated and SMR-treated samples. The strain used to prepare the cycloheximide-induced colliding ribosomes also expressed the NGD-CGA reporter, so we anticipated that within our population of bulk colliding ribosomes would be a sub-population of colliding ribosomes on our reporter, all subject to cleavage by the SMR domain. We performed RNA-seq by standard means from the trisomes treated with or without the Cue2 SMR domain and aligned the 3’ ends of the RNAs to the reporter sequence. In the sample treated with the wild type Cue2-SMR domain, we see a striking accumulation of 3’ fragment ends that map precisely where the strong cleavage sites were reported (in the A site of the colliding ribosome) in the ribosome profiling data for the same strain (*slh1Δ dom34Δ sk2iΔ*); reassuringly, these cleavage events are not seen in the sequencing data for the Cue2 SMR-R402A mutant or for the no treatment control (Figure 5B). The remarkable agreement between the *in vivo*-based ribosome profiling data and the *in vitro*-based cleavage data provides strong support for the identity of Cue2 as the endonuclease involved in NGD.

### Ribosome profiling provides evidence that Slh1 inhibits ribosome accumulation on problematic mRNAs

Our analyses (northern blots and profiling) provide strong evidence that deletion of *SLH1* in yeast results in increased levels of Cue2 cleavage on our NGD-CGA reporter (as well as on endogenous NGD mRNAs). Multiple molecular mechanisms could be responsible for this outcome. For example, a simple model might involve direct competition on the colliding ribosomes wherein Slh1 binding simply blocks Cue2 from binding to the same site. Another model could be that Slh1 functions to clear colliding ribosomes, dissociating large and small subunits from one another, and thus allowing the released peptidyl-tRNA:60S complex to be targeted for RQC (Matsuo). Other studies have argued that Slh1 functions as a translational repressor though the dependence of RQC activity on Slh1 function argues against this model (Searfoss, Dever, & Wickner, 2001).

We used ribosome profiling to directly assess ribosome occupancy on our problematic mRNA sequence as a function of Slh1 activity. As described above, when we compared 16 nt RPF accumulation on the NGD-CGA reporter in a *ski2Δ* or *slh1Δ ski2Δ* strain (Fig. 4D) we saw no differential accumulation of these RPFs that might be indicative of Slh1 function on ribosomes trapped on truncated mRNAs. However, when we compared RPFs on the uncleaved mRNAs (i.e. the 21 and 28 nt RPFs) in the *cue2Δ dom34Δ ski2Δ* and *slh1Δ cue2Δ dom34Δ ski2Δ* strains, we observed that *SLH1* deletion results in a substantial build-up of RPFs both at the stalling site (21 nt RPFs) (compare middle panels, Fig. 4C and 4F) and upstream (28 nt RPFs) of the stalling site (compare bottom panels, Fig. 4C and 4F and quantitation in 4H). These data were normalized to overall mRNA levels and reveal more than a 2-fold increase in ribosome occupancy when *SLH1* is deleted. These data are consistent with the proposed model that Slh1 prevents the accumulation of colliding ribosomes (Matsuo et al., 2017) thereby limiting the availability of Cue2 substrates.

### Genome-wide exploration of endogenous mRNA substrates of Cue2 and Slh1

One possible endogenous target class for Cue2 is the set of prematurely polyadenylated mRNAs (where polyadenylation happens within the annotated ORF) that are commonly generated by aberrant RNA processing events in the nucleus. We looked for such candidates in our data sets by first identifying genes that exhibit evidence of untemplated A’s within the monosome footprint population (i.e. ribosomes that are actively translating prematurely polyadenylated mRNAs) (Fig. 6A, pink dots). These candidate genes nicely correlated with those previously identified from 3’ end sequencing approaches (Ozsolak et al., 2010; Pelechano, Wei, & Steinmetz, 2013) (e.g. see data for two specific mRNAs *RNA14* and *YAP1* in Fig. S6A) and from ribosome profiling approaches exploring Dom34 function (N. Guydosh & Green, 2017). We next identify Cue2-dependent 16 nt RPFs in both a *ski2Δ dom34Δ* (Fig. 6A, orange dots) as well as in a *ski2Δ dom34Δ slh1Δ* background (Fig. S6B, orange dots) and see that they are substantially enriched in the prematurely polyadenylated mRNAs (Fig 6A and 6SB, red dots). We note that in the *ski2Δ dom34Δ* background a good fraction (21/55) of the prematurely polyadenylated genes are substantially reduced in 16 nt RPFs upon *CUE2* deletion (Fig. 6B). Moreover, we show that for well-defined mRNA targets *RNA14* and *YAP1*, the Cue2-dependent 16 nt RPFs are positioned as anticipated directly upstream of the previously characterized sites of premature polyadenylation (Fig. 6C and Fig. S6C) (N. Guydosh & Green, 2017; Pelechano et al., 2013). These data support a broad role for Cue2-mediated endonucleolytic cleavage on such problematic transcripts in rapidly growing yeast.

**Fig. 6:**
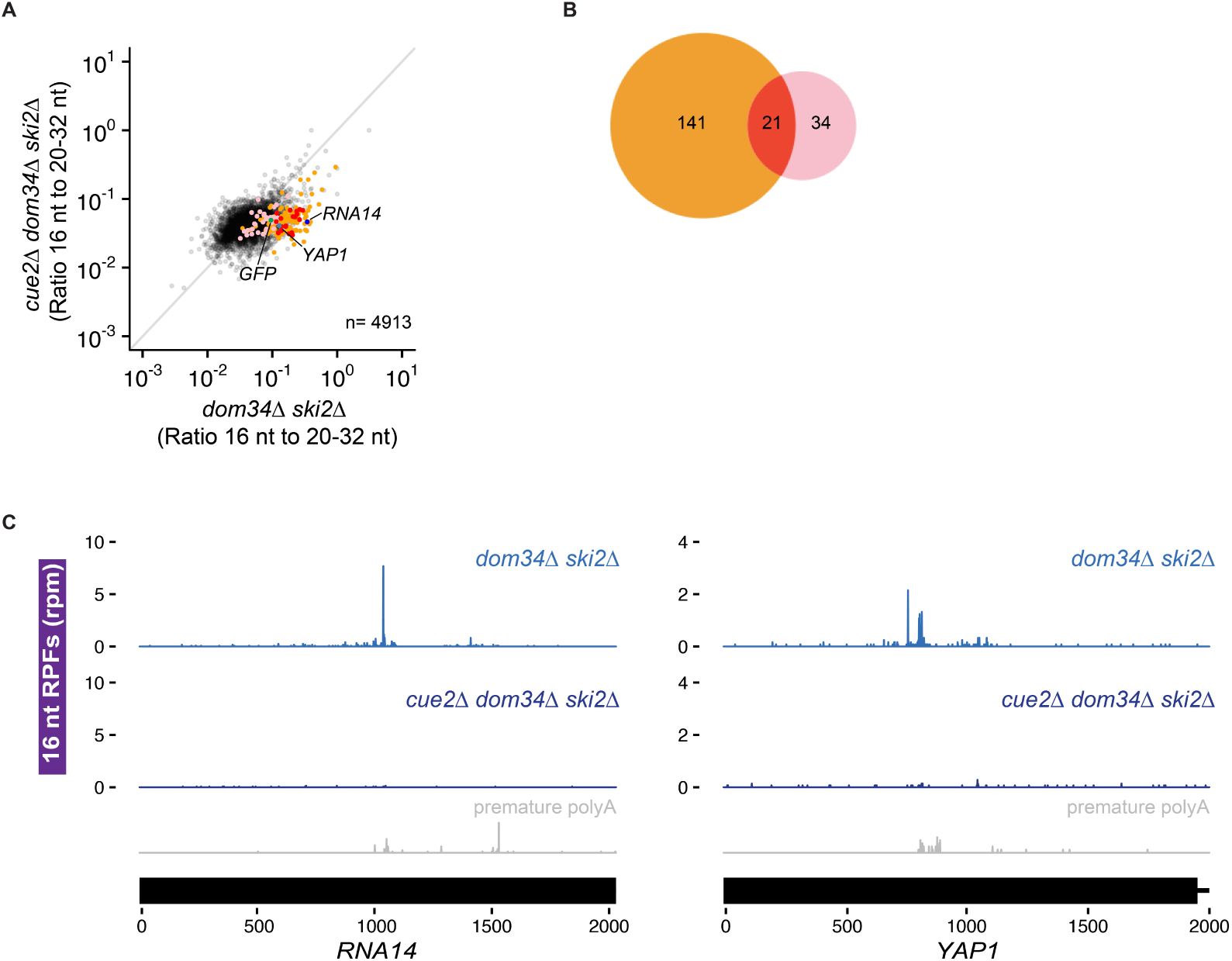
Cue2 targets prematurely polyadenylated mRNAs genome-wide for NGD. **(A)** Ratio of 16 nt over 20-32 nt RPFs plotted in *dom34Δ ski2Δ* and *cue2Δ dom34Δ ski2Δ* strains. Genes in orange correspond to those with reproducibly decreased 16 nt RPFs upon *CUE2* deletion. Pink dots indicate genes where premature polyadenylation was identified empirically from sequencing data. NGD-CGA reporter, *YAP1* and *RNA14* are labeled and shown in green, blue and purple, respectively. **(B)** Overlap (red) between annotated prematurely polyadenylated genes (pink) and genes on which CUE2 deletion had a substantial effect on 16 nt RPFs (orange). **(C)** Examples of 16 nt RPFs mapped to genes, *RNA14* (left) and *YAP1* (right), in *dom34Δ ski2Δ* and *cue2Δ dom34Δ ski2Δ* strains with known premature polyadenylation sites indicated (grey traces, from Pelechano et al., 2013 (Pelechano et al., 2013)).

## Discussion

Cells have evolved a complex set of mechanisms to recognize and resolve ribosomes stalled on problematic mRNAs and to target the mRNAs for decay. The data presented here identify and characterize Cue2 as the endonuclease critical for NGD in yeast. We demonstrate using a wide range of *in vivo* and *in vitro* approaches that Cue2 is necessary and sufficient for cleavage of mRNAs loaded with stalled, colliding ribosomes. Structural homology modeling with the C-terminal domain of IF3 positions the catalytic SMR domain of Cue2 in the A site of the small ribosome subunit (Fig. 2C and 2D, Fig S2D to Fig. S2F). And, according to this model, the conserved amino acids in Cue2 that closely approach mRNA in the A site (Fig. 2D, Fig. S2B and S2C) are indeed critical for efficient endonuclease cleavage activity (Fig. 2E and 2H). Although mutating any one of the residues D348, H350, or R402, did not completely abolish Cue2 activity, mutating a combination of these residues or removing the entire SMR domain from the protein had a very strong effect (Fig. 2E and 2H). Other studies have argued that the NGD endonuclease leaves a cyclic 3’ phosphate and a 5’ hydroxyl, products that are consistent with Cue2 being a metal independent endonuclease (Navickas et al., 2018). As metal independent RNases typically depend on mechanisms where RNA is contorted to act upon itself, it is often difficult to identify a single essential catalytic residue. Interestingly, N4BP2, the mammalian homolog of Cue2, contains a PNK domain; this kinase activity could be required for the further degradation of the mRNA through Xrn1, which depends on 5’ phosphate groups.

Strong support for the modeling of Cue2 in the A site came from the high-resolution ribosome profiling data establishing that Cue2 cleaves mRNAs precisely in the A site of the ‘colliding’ ribosome (Fig. 4). We additionally find by profiling that the lead ribosome is stalled in an unrotated state (21 nt RPFs), unable to effectively accommodate the 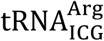 that should normally decode the CGA codon in the A site in concordance with previous studies on problematic codon pairs (Wu et al., 2019). We also find that the lagging ribosomes are found in a pre-translocation or rotated state (28 nt RPFs). These different conformational states of the colliding ribosomes identified by ribosome profiling correlate with those defined in recent cryoEM structures of colliding ribosomes (REFS). We note that our data and existing structures suggest that Cue2 requires partial displacement of the peptidyl-tRNA substrate positioned in the A site of the colliding ribosome in order to access the mRNA. Recent studies identify residues in Rps3 at the mRNA entry tunnel that are required for NGD cleavage consistent with the idea that Cue2 binds within the ribosome mRNA channel to promote cleavage (Simms et al., 2018).

Another structural feature of Cue2 is that it possesses two CUE domains at the N-terminus connected by a long linker region to the proposed catalytic SMR domain. Endonucleolytic cleavage of problematic mRNAs was previously shown to depend on the function of the E3 ligase Hel2 which ubiquitinates several small subunit ribosome proteins (Ikeuchi et al., 2019); our ribosome profiling data provides strong support for this model (Fig. S4). We suspect that the two CUE domains might recognize Hel2-ubiquitinated sites on the colliding ribosomes thus allowing recruitment of the SMR domain into the A site of the colliding ribosome. We can imagine that recognition of ubiquitin chains by Cue2 might involve multiple ubiquitinated sites on a single collided ribosome or those found on neighboring ribosomes. These CUE domains might also function within Cue2 itself to inhibit promiscuous activity of the endonuclease in the absence of an appropriate target; this possibility prompted us to perform the *in vitro* cleavage experiments with the isolated SMR domain of Cue2.

In addition to the identification and characterization of the key endonuclease involved in NGD, our results help to clarify the relative contributions of decay pathways for problematic mRNAs. We show that canonical exonucleolytic processing of problematic mRNAs is the dominant pathway on our NGD-CGA reporter, with the strongest contributions from Xrn1 rather than the exosome (Fig. 3). Importantly, we find that the Cue2-mediated endonucleolytic pathway is activated in genetic backgrounds lacking the helicase Slh1 (a member of the previously identified RQT complex) (Fig. 2F-2G). These ideas are brought together in the model in Figure 7 that outlines how multiple decay pathways converge to bring about the decay of problematic mRNAs. We anticipate that mRNAs with different problematic features might differentially depend on these pathways as suggested by previous studies (Tsuboi et al., 2012). Importantly, however, our identification of the endonuclease for NGD will allow subsequent studies to better disentangle the distinct contributions from multiple decay machineries in the cell. A challenge moving forward will be identifying how problematic mRNAs with colliding ribosomes signal to decapping factors and Xrn1 to initiate degradation (indicated with question marks in the model in Fig.7).

**Fig. 7:**
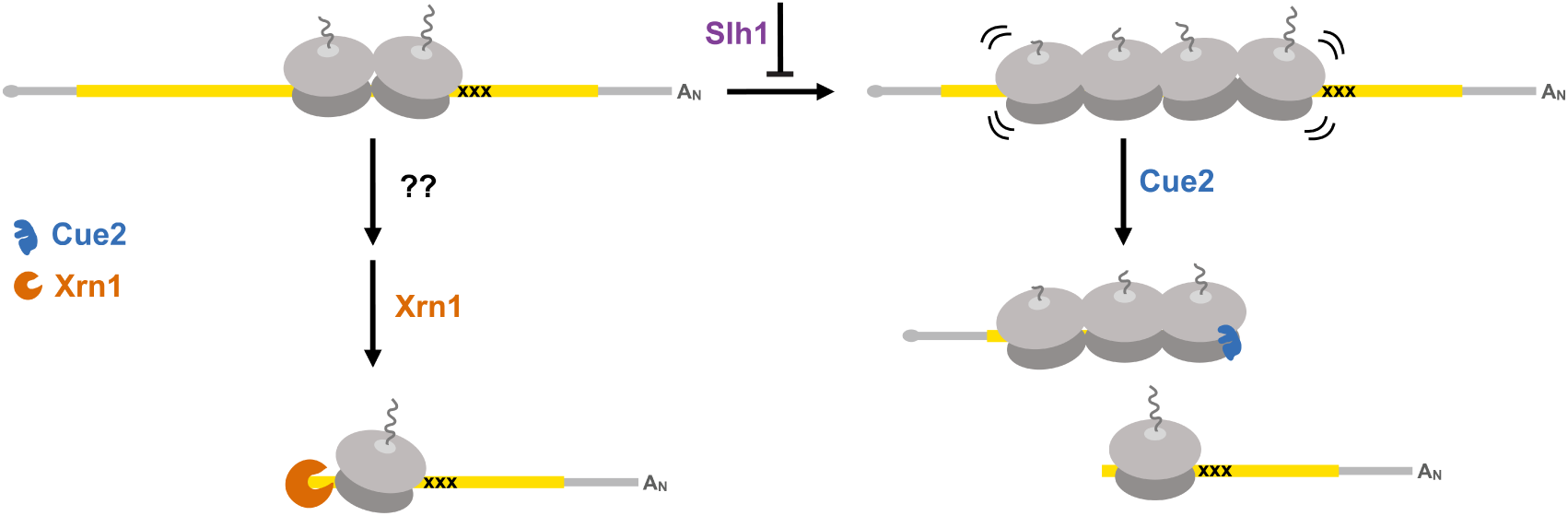
Multiple converging pathways on NGD substrates. Ribosome stalling triggers decay either by the exonuclease Xrn1 or by the endonuclease Cue2. The relative contributions of these pathways are modulated by the activity of Slh1 (member of the RQT complex). Ribosomes are grey; open reading frame is yellow; ‘xxx’ indicates a stalling sequence in the mRNA.

Our genetic insights also lead us to speculate that under conditions where cells are overwhelmed by colliding ribosome complexes, for example under stressful RNA-damaging conditions (Simms, Hudson, Mosior, Rangwala, & Zaher, 2014), that the RQT machinery might become limiting and that there will be an increased role for Cue2-mediated endonucleolytic processing. Previous genetic screens reveal increased sensitivity of *CUE2* deletion strains to ribosome-targeted compounds (Alamgir, Erukova, Jessulat, Azizi, & Golshani, 2010; Mircus et al., 2015). More generally, the strong conservation of Cue2 throughout eukaryotes argues that this endonucleolytic processing pathway plays a fundamental role in biology.

Previous studies provide strong evidence that the activity of the RQT complex is critical in targeting nascent peptides on problematic mRNAs for degradation via the RQC (Matsuo et al., 2017; Sitron et al., 2017) through ribosome rescue events that generate a dissociated 60S-peptidyl-tRNA complex. Our ribosome profiling results indicate that ribosome occupancies on problematic mRNAs increase on deletion of *SLH1* (Fig. 4C, 4F, 4H) consistent with the possibility that Slh1 functions directly or indirectly to dissociate ribosomes. The increased CUE2-dependent cleavage (Fig. 2F-2G and Fig. 4B, 4E) that we observe in *slh1Δ* background suggests that the increase in the density of collided ribosomes makes an ideal Cue2 substrate.

Taken together, we suggest that Slh1 and Dom34 may both target collided ribosomes for the RQC, though their specificities may be somewhat distinct. For example, Slh1 may use its helicase function to target ribosomes on intact mRNAs with an accessible 3’ end, while Dom34 targets those ribosomes on truncated mRNAs that are inaccessible to Slh1 activity. In this view, a critical role of Cue2-mediated endonucleolytic cleavage is to provide another pathway for ribosome dissociation that can target nascent peptides for RQC. Previous studies provide strong support for the idea that Dom34 preferentially rescues ribosomes that are trapped on truncated mRNAs (N. R. Guydosh & Green, 2014; Shoemaker et al., 2010) and *in vitro* studies have nicely connected this rescue activity of Dom34 to downstream RQC functions (Shao, Von der Malsburg, & Hegde, 2013). These two distinct rescue pathways, one on intact mRNA by the RQT complex (and Slh1) and the other on endonucleolytically cleaved mRNA by Dom34, might provide the cell with redundant means to deal with toxic mRNP intermediates.

Proteotoxic stress is a critical problem for all cells and elaborate quality control systems have evolved to minimize its effects as made clear by the exquisite sensitivity of neurons to defects in these pathways (Bengtson & Joazeiro, 2010; Ishimura et al., 2014). The synergistic contributions of exonucleases and endonucleases that we delineate here for targeting problematic mRNAs for decay are critical for ensuring that proteotoxic stress is minimized. Future studies will delineate the biological targets and conditions in which these systems function.

## Supporting information

Supplemental Table S2_strains

Supplemental Table S1_primers and plasmids

## Acknowledgments

We thank Miguel Pacheco for help with yeast experiments during the crunch. We thank Brendan P. Cormack for helpful consultation on ideas and techniques throughout this study, Ryan McQuillen for help with figures, and members of the Greider lab, Carla Connelley, Rebecca Keener, and Calla Shubin, for assistance with techniques and careful interpretations of yeast genetic experiments. We thank Allen R. Buskirk, Boris Zinshteyn, Daniel Goldman, Kamena K. Kostova, Chirag Vasavda, and Jamie Wangen for careful reading of the manuscript and all Green lab members for helpful discussions throughout this study.

## Funding

This work was supported by the NIH (R37GM059425 to R.G. and 5T32GM007445-39 training grant for K.N.D.), HHMI (to R.G.), and the Canadian Institutes of Health Research (FDN-159913 to G.W.B.).

## Author contributions

K.N.D. and R.G. conceived this study. K.N.D. constructed all strains and plasmids, and performed experiments unless otherwise specified here. C.C.W. performed all ribosome profiling experiments with all relevant data analysis. N.S. provided structural homology modeling. R.L-K. assisted in screen-related data acquisition and curation. K.N.D., C.C.W., R.L-K., G.W.B., and R.G. were responsible for methodology. K.N.D. C.C.W. and R.G. wrote the original draft of this manuscript and all authors revised and edited. Funding and resources provided by R.G. and G.W.B.

## Competing interests

Authors declare no competing interests.

## Data and materials availability

Raw sequencing data were deposited in the GEO database under the accession number GSE129128. Secure token for reviewers: snupcguclhifhex.

## Supplementary Materials

### Materials and Methods

(Armougom et al., 2006; Biou et al., 1995; Diercks et al., 2008; Dobin et al., 2013; Douglas et al., 2012; Gamble, Brule, Dean, Fields, & Grayhack, 2016; Geer, Domrachev, Lipman, & Bryant, 2002; Giaever et al., 2002; Goddard, Huang, & Ferrin, 2007; Hendry, Tan, Ou, Boone, & Brown, 2015; Y. Hu et al., 2007; Hussain et al., 2016; Jiang, Lei, Ding, & Zhu, 2014; Kainth et al., 2009; Kelley, Mezulis, Yates, Wass, & Sternberg, 2015; Longtine et al., 1998; Marchler-Bauer et al., 2011; Pettersen et al., 2004; Radhakrishnan et al., 2016; Robert & Gouet, 2014; Tong & Boone, 2006; Wagih et al., 2013; Waterhouse et al., 2018; Wu et al., 2019)

