## Supplemental Figures and Methods for "The endonuclease Cue2 cleaves mRNAs at stalled ribosomes during No Go Decay"

**This PDF file includes:**

Materials and Methods

Figs. S1 to S6

**Other Supplementary Materials for this manuscript include the following:**

Tables S1 to S2 (as .xlsx)

Materials and Methods

**Plasmid construction**

*GFP reporters*

The OPT reporter plasmid, or pKD065, was cloned in pECB1806 (*50*) as follows. Briefly, the *GAL1* promoter, a *GFP-2A-FLAG*, a partially non-optimal *HIS3* and the *ADH1* terminator were introduced in *Spe*I/*Sph*I- digested pECB1806 using Gibson assembly leading to pKD064. The *GAL1* promoter was amplified by PCR using primers KD235 and KD236 and the *ADH1* terminator was PCR amplified using KD239 and KD240 with pECB1806 as template (Table S2). All PCR products were cleaned using Zymo Research DNA Clean and Concentrator Kit. The *GFP-2A-FLAG* and a partially non-optimal *HIS3* sequence fragment was gene-synthesized by Integrated DNA Technologies (IDT). The partially non-optimal *FLAG-HIS3* in pKD064 was then replaced by a fully optimal sequence of *FLAG-HIS3* from pJC797 (*64*) to make pKD065 (Table S2).

The NGD-AAA and NGD-CGA reporter plasmids (pKD079 and pKD080, respectively) were cloned using pKD065. In brief, a (AAA)x12 or (CGA)x12 codon sequence was inserted 90 codons into the *HIS3* gene of pKD065. The PCR product from (AAA)12 primers (KD281 and KD280) or (CGA)12 primers (KD283 and KD280) with pKD065 template was combined with the PCR product from primers KD287 and KD276 with pKD065 template and *Sal*I/*Sph*1 digested pKD065 and these products were Gibson assembled to produce pKD079 and pKD080 (Table S2).

*HA-CUE2 overexpressing plasmid*

The CUE2 overexpressing FLEX plasmid (pKD100) was rescued from the yeast FLEX library (*33, 55*). A gene block (gKD002) of truncated CUE2 (NoSMR) was inserted into *BamH*I/*Sph*I digested pKD100, to make pKD105. 5’ 3xHA-tagged *CUE2*, and *NoSMR* were PCR amplified using KD414 and KD416 and Gibson assembled with *BamH*I/*Sph*I digested pKD100 to make pKD120 and pKD125, respectively (Table S2).

*Site-directed mutagenesis*

Using the standard protocol for the QuikChange Lightning Multi Site-Directed Mutagenesis Kit from Agilent Technologies, the indicated mutations in the *HA-CUE2* construct (pKD120) were made to make pKD127, pKD129, pKD131, pKD133, and pKD145 (Table S2).

*CRISPR plasmids*

BplI digested plasmid pJH2972 (Anand, Memisoglu, & Haber, n.d.) was used in a Gibson reaction with primers KD503, KD505, and KD509 to make plasmids pKD163, pKD165, and pKD169 respectively.

*Cue2 E. coli expression plasmids*

PCR products from KD457 and KD338 from plasmids pKD120 and pKD129 were Gibson cloned into BamHI and XhoI digested pSMT3, to make pKD097 and pKD098, respectively.

**Yeast Strains and growth conditions**

Yeast strains used in this study are described in Table S2 and are all derivatives of BY4741 unless specified otherwise. Yeast strain construction was performed using standard lithium acetate transformations. Reporters strains were constructed by integrating the various GFP-2A-FLAG-HIS3 cassettes, from *Stu*I digested pKD065, pKD079, and pKD080, into the *ADE2* locus of BY4741 (Table S2). For SGA experiments, query strains overexpression screens were constructed by introducing the GFP-2A-FLAG-HIS3 cassettes from *Stu*I digested pKD065, pKD079, or pKD080 at the *ADE2* locus in BY4741 (Table S2).

Deletion strains were constructed by inserting resistance cassettes from Longtine et al. 1998 (*57*) into the designated loci and genotypes are listed in Table S2. Note: two different deletion strains were used for *XRN1* deletions in this study. See Table S2 for genotypes.

HA tag insertions, point mutations, and the SMR deletion were made in yKD143 as described in Anand et al. 2017, using plasmids pKD163, pKD165, and pKD169 and homology directed repair templates.

Recipes for media used in this study are listed in Table S2.

For gal-induced growths, overnight cultures were grown in YPAGR media, or, for strains with plasmids, were grown in SC/A/GR/-Ura. Overnights were diluted in the same media to an OD of 0.1 and harvested at an OD of 0.4-0.5.

**Flow cytometry:**

*Data Collection:*

100 μl of log-phase cells were pelleted and washed once with 1 x PBS. Cells were then resuspended in 500 μl of 1 x PBS and 5000 cells were analyzed with a Millipore guava easyCyte flow cytometer for GFP and RFP detection using 488 nm and 532 nm excitation lasers, respectively.

*Data Analysis:*

Cells were gated based on size, and random outliers were cut off from graphs for visual purposes (plot values were not changed upon removal of outliers). For all flow cytometry data, violin plots show the density of cells for log_2_(GFP/RFP) values for individual cells.

**Northern Blots:**

*RNA isolation*

25 mls of log-phase cells were pelleted and supernatant was poured off. Cells were resuspended in residual media, pelleted again, and flash frozen in liquid nitrogen. Cell pellet was resuspended in buffer with 8.4 mM EDTA, and 60 mM NaOAc. 20% SDS was added to a final concentration of 1.5%. Cell solution was warmed at 65°C for 2 minutes and added to acid phenol at 65°C. Phenol/cell solution was shaken at 1100 rpm at 65°C for 10-20 minutes with intermittent vortexing. Samples were put on ice for 5 minutes then spun at 16 krpm. The aqueous layer was removed and added to an equal volume of phenol, and samples were vortexed again. Samples were spun at 16 krpm again, the aqueous layer was removed and added to an equal volume of chloroform, and samples were vortexed. Lastly, samples were spun again at 16 krpm, the aqueous layer was removed and precipitated in NaOAc and isopropanol. The RNA pellet was resuspended in 10 mM Tris-HCl, pH 8.0.

*Northern blot*

Between 5 and 10 μg of RNA were loaded into a 1.2% agarose, formaldehyde denaturing gel and run for 2-2.5 hours at 125 volts (for any given gel, the same amount of RNA was loaded for each sample, but this varied from gel to gel). The RNA was vacuum transferred to a nitrocellulose (N+ Hbond, Amersham) membrane in 10 x SSC buffer. The RNA was then UV crosslinked to the membrane and placed in pre-hybridization buffer and rotated at 42°C. The indicated PNK-end labeled probe was added to the pre-hybridization buffer at 42°C after 30 minutes (Table S2). The membrane was probed overnight, rotating at 42°C. The membrane was washed 3 times in 2 x SSC, 0.1% SDS for 20 minutes at 30°C, then exposed to a phosphoscreen. The phosphoscreen was scanned using a Typhoon FLA 9500.

*End-labeled DNA probe*

This indicated DNA-oligo listed in Table S2 was end labeled using gamma-ATP and the standard T4 Polynucleotide Kinase radioactive labeling protocol from NEB. The labeled oligo was purified using GE Healthcare illustra ProbeQuant G-50 Micro Columns.

**Western Blots:**

*Protein Isolation*

2 OD units of log-phase cells were pelleted and supernatant was poured off. Cells were resuspended in residual media, pelleted again, and flash frozen in liquid nitrogen. Pellets were resuspended in 200 μl lysis buffer containing 20mM Tris-HCl, pH 8.0, 140mM KCl, 5mM MgCl2, 1% triton, 1mM DTT, Roche cOmplete protease inhibitor tablet, and PMSF, pepstatin, and leupeptin protease inhibitors. The volume of cell solution was approximately doubled using acid washed glass beads and cells were mechanically lysed using bead beater 1 min on, 1 min off, for 3 cycles. 6x SDS loading dye was added to the lysis solution at 2x and samples were boiled for 5 min.

*Western blot*

Equal volumes of lysate were loaded for each sample onto 4-12% Criterion XT Bis-Tris protein gels in 1x XT MES buffer. Protein was transferred to PVDF membrane via turbo blot. Membranes were then placed in 2.5% milk, 1x TBST blocking solution for 1 hour. Primary antibody was used in 1x TBST at 1:5000 for anti-PGK1 (Invitrogen, mouse), anti-HA (Roche, rat), anti-FLAG (Sigma, mouse), or anti-GFP (Takara, mouse) and incubated on a rotator overnight at 4°C. Membranes were washed in 1x TBST 3 times, 10 minutes each. The corresponding HRP-conjugated secondary antibody was added to the membrane at 1:5000 in 1x TBST and incubated for 1-2 hours. Membranes were washed 3 times for 10 minutes each. Pico solution (details) was added to membranes for approximately 3 minutes and then membranes were scanned using a G:BOX Chemi XX6 (Syngene) with varying exposure times.

**Reporter-SGA screens**

*SGA procedure*

SGA screens were performed using a Biomatrix Robot (S&P Robotics Inc.) with a few modifications (*61*). Briefly, yKD131, yKD132, yKD133 query strains (Table S2) were crossed individually with the FLEX collection (*32*, *33*). The FLEX library was arrayed in a 1536-format containing 4 colonies for each overexpression strain. Mating steps were performed on standard SGA media (*61*).

For the overexpression screen, diploid strains were selected on OEDIP media listed in Table S2. Sporulation and haploid double mutant selection steps were not performed in this screen.

To induce reporter expression and FLEX gene overexpression, cells were pinned again onto the same medium (diploid selection medium for the FLEX screen) except that glucose was replaced by raffinose and galactose at a final concentration of 2% (OEDIPGR media listed in Table S2). Cells were grown for 40-46 hours for the overexpression screen before scanning on a Typhoon FLA9500 (GE Healthcare) fluorescence scanner equipped with 488 nm and 532 nm excitation lasers and 520/40 and 610/30 emission filters. Plates were also photographed using a robotic system developed by S&P Robotics Inc. in order to determine colony size.

*Screen data analysis:*

GFP and RFP fluorescent intensity data was collected using the microarray software, TIGR Spotfinder (Saeed AI, Sharov V, White J, Li J, Liang W, et al. TM4: a free, open-source system for microarray data management and analysis. Biotechniques. 2003;34:374–378). Colony size data was aquired using SGATools (*62*) (<http://sgatools.ccbr.utoronto.ca/>). Subsequent data analysis was performed as previously described (*53*, *55*). In brief, border strains and size outliers (<1500 or >6000 pixels) were eliminated, and median GFP and RFP values were taken for the remaining strains. Log_2_(mean GFP/mean RFP) values were then calculated and LOESS normalized for each plate. Finally, Z-scores for each individual plate were calculated based on the LOESS normalized log_2_(mean GFP/mean RFP). Strains for the AAA or CGA reporters with Z-scores greater than 2.5 or less than -2.5 were considered as hits if their Z-score in the OPT reporter was unaffected (i.e had a Z-score between -2.5 and 2.5). For subsequent experiments, hits were reconstructed in the background strain yKD133 as described above and all experiments were performed with the reconstructed strain.

*Venn Diagrams:*

Overexpression screen strains with a Z-score greater than 2.5 or less than -2.5 were considered candidate genes and analyzed for overlap using BioVenn (<http://www.biovenn.nl/>) to produce the diagrams.

**Cue2-SMR prep**

The SMR and the R402A-SMR constructs, pKD097 and pKD098, respectively, were expressed in RIPL BL21 E. coli strain using Kan resistance and inducing at OD=0.4, at 18°C with IPTG at 0.5 mM overnight. Cells were harvested by centrifugation and then flash frozen in liquid nitrogen. Pellets were then resuspended in protein prep lysis buffer [25 mM Tris-Cl pH 7.5, 500 mM KCl, 1 mM MgCl2, 5 mM bMe, 10% glycerol] + PMSF, leupeptin, pepstatin, and Roche cOmplete EDTA-free protease inhibitors.

Cells were lysed using the French press, 3 times, 1100 pressure units. Lysis solution was clarified at 20000xg for 30min. Lysate was filtered through a 0.2 um filter and run over HiTrap 5ml Ni-NTA column. Column was washed for 5 CV in lysis buffer with high salt (1 M KCl) and 20 mM imidazole. Sample was batch eluted using Ni elution buffer [25 mM Tris-Cl pH 7.5, 500 mM KCl, 1 mM MgCl2, 5 mM bMe, 10% glycerol, 500 mM imidazole]. Sample was diluted 10 fold into S column buffer [25 mM Tris-Cl pH 7.5, 200 mM KCl, 5 mM MgCl2, 5 mM bMe, 10% glycerol] and then run on a Resource S column. Sample was washed in S column buffer for 5 CV and then gradient eluted off of S column in S column buffer with 1 M salt. 6xHIS-Sumo tag was cleaved overnight using SUMO protease. And sample was run on an orthogonal Ni column to remove tag. Flow through was collected and concentrated to 40 uM SMR, and 20 uM SMR-R402A.

**Isolation of nuclease resistant trisomes**

Grow 1 liter of yKD307 in YPAGR to an OD 0.4. For cycloheximide treated cells, added 1 mg/L cycloheximide, grew for 30 minutes and filter harvested. For untreated cells, immediately filter harvested. Cell pellets were ground with 1 mL lysis buffer [10 mM potassium phosphate (pH 6.1), 140 mM KCl, 5 mM MgCl_2_, 1% Triton X-100, 0.1 mg/mL cycloheximide, 1mM DTT, Roche cOmplete EDTA-free protease inhibitor, leupeptin, PMSF, pepstatin] in a Spex 6870 freezer mill. Cell lysates were clarified by centrifugation. CaCl2 was added to 200 OD units of clarified lysates to a final concentration of 2.5 mM and the indicated clarified lysates were treated with 20 ul of NEB MNase at 2e^6 gel units/ml for 30 minutes. 4 mM EGTA was added to quench the reaction. Samples were layered on a 15-45% sucrose gradient [10 mM potassium phosphate (pH 6.1), 140 mM KCl, 5 mM MgCl_2_] and spun in SW28 rotor at max speed for 4 hours. Nuclease resistant trisome peak was isolated and diluted two fold in lysis buffer.

**Cleavage of lysates with SMR and analysis**

Protein prep buffer, the purified SMR domain of Cue2, and the R402A mutant of the SMR domain of Cue2 were added to separate aliquots of isolated nuclease resistant trisomes, such that the final concentration of enzyme (for samples with enzyme) = 3 µM. Samples were left at room temperature for 2 hours. Next, samples were run on a 15-45% sucrose gradient [10 mM potassium phosphate (pH 6.1), 140 mM KCl, 5 mM MgCl_2_] and the A260 trace is reported. RPFs were extracted from pooled fractions (mono-, di-, and trisomes).

**Sequence alignment**

Information from NCBI conserved domain database (*58*), conserved domain architecture retrieval tool (*51*), available structures, and Phyre-based homology modeling (*56*) was used to define domain boundaries. Structure-based multiple sequence alignments were carried out using Expresso (*48*), and illustrated with ESPript (*60*). Protein names are indicated, followed by the name of organism, and residues used for alignment. YEAST, *Saccharomyces cerevisiae*; CANGA, *Candida glabrata*; ARATH, *Arabidopsis thaliana*; HUMAN, *Homo sapiens*; MACMU, *Macaca mulatta*; BOVIN, *Bos taurus*; THETH, *Thermus thermophilus*; BACSU, *Bacillus subtilis*; ECOLI, *Escherichia coli*.

**Homology modeling**

Homology model of the Cue2 SMR domain templated on human N4BP2 SMR domain (PDB: 2VKC (*38*) was generated using Phyre 2.0 (*56*) and SWISS-MODEL (*63*). After identifying N4BP2 as a Cue2 homolog, a heuristic DALI search was performed using the N4BP2 SMR domain (PDB IDs: 2D9I and 2VKC) as query against structures in the Protein Data Bank (PDB). This exercise, along with templates identified using Phyre 2.0 (*56*) and SWISS-MODEL (*63*), enabled us to identify structural conservation between the SMR domain of Cue2 and the C-terminal domain (CTD) of translational initiation factor 3 (IF3). This was further validated using structure-based sequence alignments as shown in Fig. S2B. Structural comparisons and fittings were performed using UCSF Chimera (*52*, *59*). Superimposition of the CTD of IF3 (PDB: 1TIG (*39*)) and the Cue2-SMR homology model revealed structural conservation between the two domains (Fig. S2C). This enabled superimposition of the Cue2 SMR in the context of full-length Thermus thermophilus IF3 (PDB: 5LMQ (*40*)) (see Fig. 2C). This fitting enabled overlay of the Cue2-SMR at the A/P-site in the context of IF3 and tRNAfMet bound 30S pre-initiation complex (PDB: 5LMQ, State 2A (*40*)) (see Fig. 2D and Fig. S2, D-E). The mRNA and the 30S ribosomal subunit are represented as surfaces to minimize over interpretation (Fig. 2D and Fig. S2, D-E).

**Yeast growth conditions for ribosome profiling**

Overnight seed cultures were grown in YPAD at 30°C. To induce NGD-CGA reporter expression, cells were collected by centrifugation, washed, and resuspended in YPAGR. Cells were then harvested at OD ~0.5 by fast filtration and flash frozen in liquid nitrogen.

**Preparation of libraries for yeast ribosome footprints**

Cell pellets were ground with 1 mL footprint lysis buffer [20 mM Tris-Cl (pH8.0), 140 mM KCl, 1.5 mM MgCl_2_, 1% Triton X-100, 0.1 mg/mL cycloheximide, 0.1 mg/mL tigecycline] in a Spex 6870 freezer mill. Lysed cell pellets were diluted to 15 mL in footprint lysis buffer and clarified by centrifugation. The supernatant was layered on a sucrose cushion [20 mM Tris-Cl (pH8.0), 150 mM KCl, 5 mM MgCl_2_, 0.5 mM DTT, 1M sucrose]. Polysomes were pelleted by centrifugation at 60,000 rpm for 106 min in a Type 70Ti rotor (Beckman Coulter). Ribosome pellets were gently resuspended in 800 µL footprint lysis buffer. 350 µg of isolated polysomes were treated with 500 units of RNaseI (Ambion) for 1 hr at 25˚C. Monosomes or disomes were isolated by sucrose gradients (10-50%). RNA was extracted by hot acid phenol and then size-selected from 15% denaturing PAGE gels, cutting between 15-34 nt for monosome footprints and 40-80 nt for disome footprints. Library construction was carried out as described (*45*). Libraries were sequenced on a HiSeq2500 machine at facilities at the Johns Hopkins Institute of Genetic Medicine.

**Analysis of ribosome profiling data**

The R64-1-1 S288C reference genome assembly (SacCer3) from the *Saccharomyces* Genome Database Project was used for yeast genome alignment. For the rest of our libraries, 3’ adapter (NNNNNNCACTCGGGCACCAAGGA) was trimmed, and 4 random nucleotides included in RT primer (RNNNAGATCGGAAGAGCGTCGTGTAGGGAAAGAGTGTAGATCTCGGTGGTCGC/iSP18/TTCAGACGTGTGCTCTTCCGATCTGTCCTTGGTGCCCGAGTG) were removed from the 5’ end of reads with skewer (*54*). Trimmed reads were aligned to yeast ribosomal and non-coding RNA sequences using STAR (*49*) with ‘--outFilterMismatchNoverLmax 0.3’. Unmapped reads were then mapped to genome using the following options ‘--outFilterIntronMotifs RemoveNoncanonicalUnannotated --outFilterMultimapNmax 1 --outFilterMismatchNoverLmax 0.1’. All other analyses were performed using software custom written in Python 2.7 and R 3.3.1.

For monosome footprints, the offset of the A site from the 5’ end of reads was calibrated using start codons of CDS (*45*). For 28 nt RPFs, offsets (27:[16], 28:[16], 29:[17], 30:[17], 31:[17], 32:[17]) were used to infer the A sites of 27-32 nt reads. For 21 nt RPFs, offsets (20:[16], 21:[17], 22:[17]) were used. For 16 nt RPFs, 3’ ends of 15- 17 nt RPFs were used to infer cleavage sites. For 60 nt disome footprints, offsets (57:[47],58:[47],59:[47],60:[47],61:[47],62:[47]) were used to infer the A sites of lead ribosomes. For 54 nt disome footprints, offsets (51:[47],52:[47],53:[47],54:[47]) were used. For 46 nt disome footprints, 3’ends of 44-48 nt RPFs were used to infer cleavage sites. Prematurely polyadenylated mRNAs were identified by monosome footprints (15-34 nt) with the criteria of at least three RPFs of more than one untemplated A’s in the 3’end from both *slh1∆ cue2∆ dom34∆ ski2∆* datasets. Cue2 target genes were identified using 15-17 nt RPFs that exhibited reproducible reduction upon *Cue2* deletion (adjusted p < 0.5, DESeq (Ref: Anders and Huber, 2010)). For *in vitro* cleavage assays, 3’ ends of 60-65 nt RPFs isolated from pooled fractions were used to infer cleavage sites (Figure 5B).

**Accession number**

Raw sequencing data were deposited in the GEO database under the accession number GSE129128. Secure token for reviewers: snupcguclhifhex.

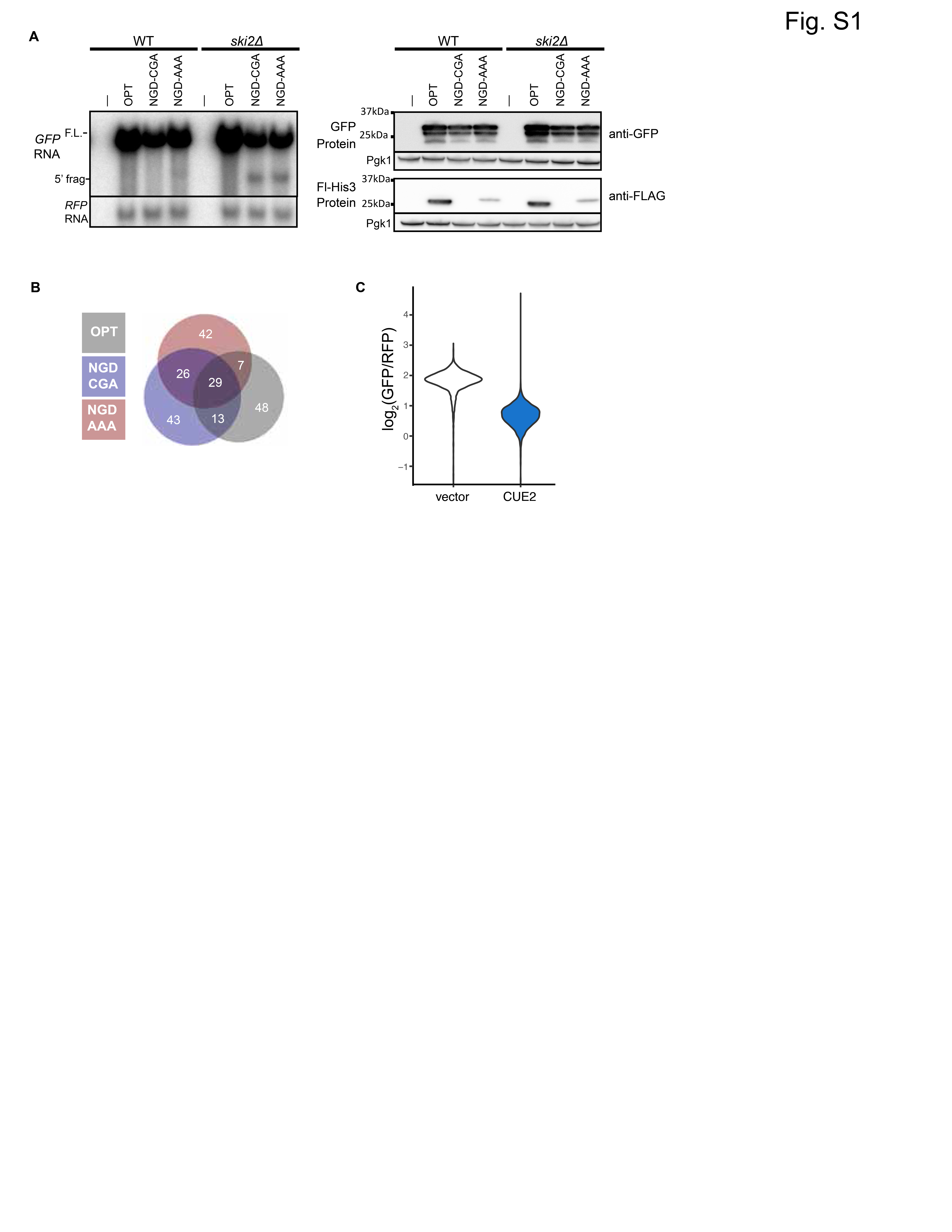

**Fig. S1. (A)** Northern and western blot analysis for the indicated strains and reporters. Right, flow cytometry data for the indicated reporters in the *ski2∆* strain. **(B)** Venn diagram of top outliers from the overexpression screen. (C) Raw flow cytometry data from >4000 cells from the empty vector and *CUE2* overexpression strains. One replicate of the triplicate in Figure 1E.

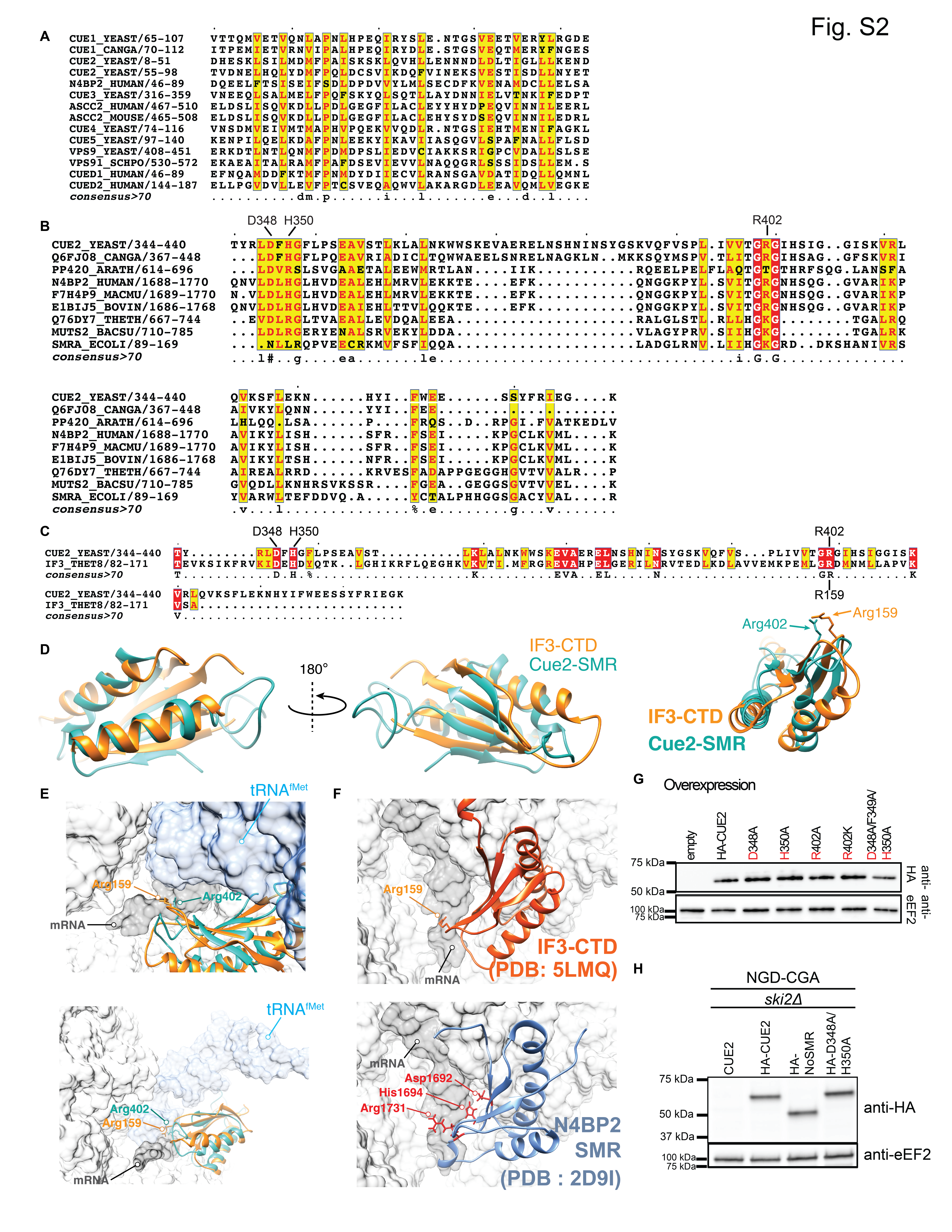

**Fig. S2. (A)** Sequence alignment of CUE domains of representative proteins. Conserved residues depicted in red on yellow background. **(B)** Sequence alignment of SMR domains of representative proteins. Identical residues shown in white on red background; conserved residues shown in red on yellow background. **(C)** Structure-based sequence alignment of the SMR domain of *Saccharomyces cerevisiae* Cue2 (residues 344-440) with the C-terminal domain (CTD) of *Thermus thermophilus* translation initiation factor 3 (IF-3) (residues 82-171). Identical residues depicted in white on red background; conserved residues depicted in red on yellow background. **(D)** Superimposition of the CTD of IF3 (orange, PDB: 1TIG (Biou, Shu, & Ramakrishnan, 1995)) and a homology model of the SMR domain of Cue2 (cyan) showing structural conservation between the two domains. **(E)** Modeling of the Cue2-SMR homology model in the context of IF3 and tRNA^fMet^ bound 30S pre-initiation complex (PIC) (PDB: 5LMQ, State 2A (Hussain, Llácer, Wimberly, Kieft, & Ramakrishnan, 2016) , also see Fig. 2C-D). Light gray, 30S; dark gray, mRNA; blue, tRNA^fMet^; orange, IF3; cyan, Cue2-SMR. Vignettes show the position of Arg-159 of IF3 near the AUG-start codon at the P site, with the putative position of the catalytic Arg-402 of Cue2-SMR in the vicinity of the mRNA, based on homology modeling. **(F)** Alternate view of IF3-CTD (top panel) bound to 30S pre-initiation complex (PIC) with the tRNA^fMet^ density subtracted (PDB: 5LMQ, State 2A, (Hussain et al., 2016)), and N4BP2 SMR domain (bottom panel) showing putative positions of the corresponding Asp-1692, His-1694 and Arg-1731 residues (PDB: 2D9I (Diercks et al., 2008)). **(G)** Western blot analysis of HA tagged WT and mutant Cue2 overexpression levels for flow cytometry data in Fig. 2E. **(H)** Western blot analysis of HA tagged endogenous WT and mutant Cue2 levels for northern blot analysis in Fig. 2H.

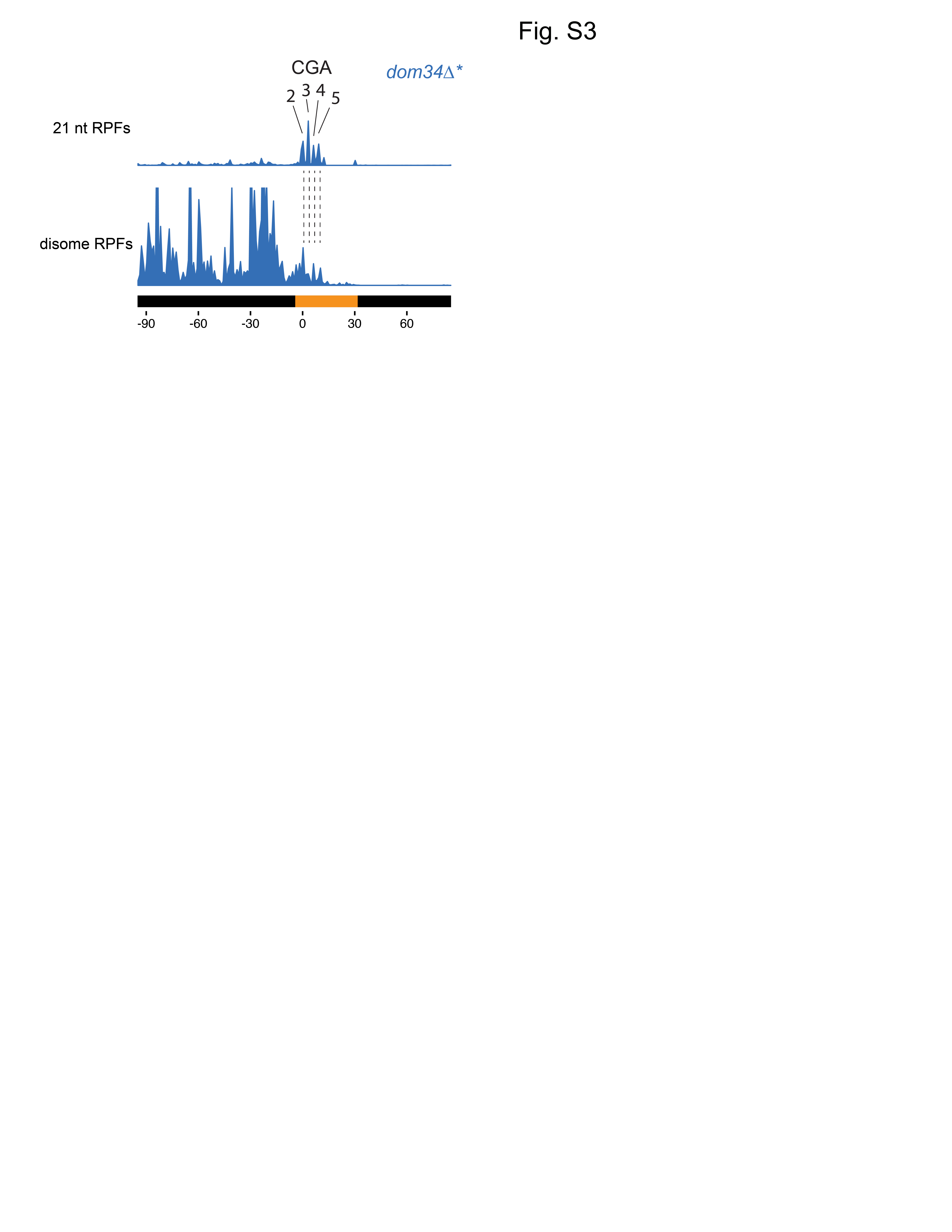

**Fig. S3:** Comparison between monosome 21 nt RPFs and disome footprints. Disome footprints show predominant ribosome stall site at the 2^nd^ to 5^th^ CGA codons.

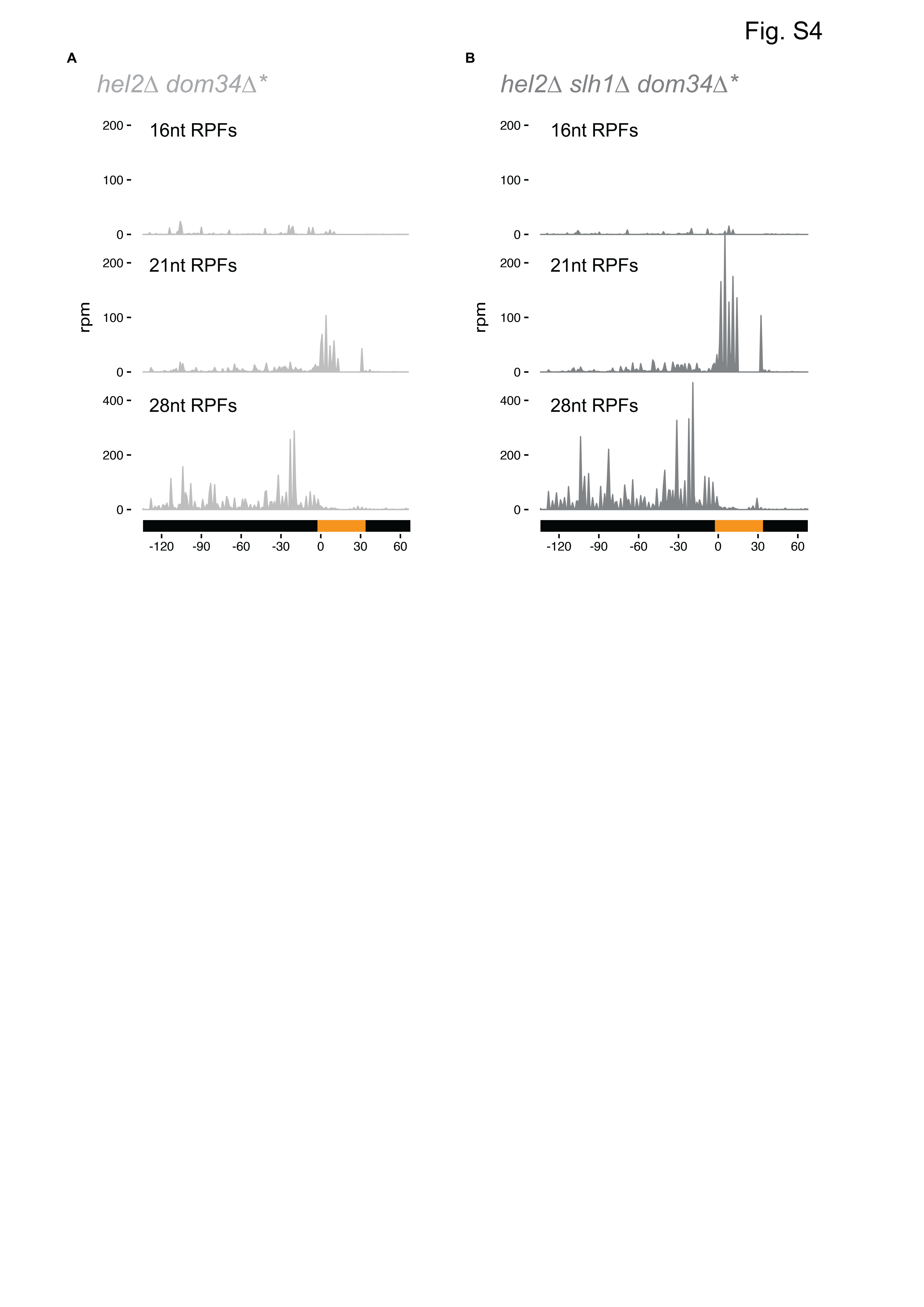

**Fig. S4:** 16, 21 and 28 nt RPFs on the NGD-CGA reporter from *hel2∆ dom34∆* *ski2∆* **(A)** and *hel2∆ slh1∆ dom34∆* *ski2∆* **(B)** strains.

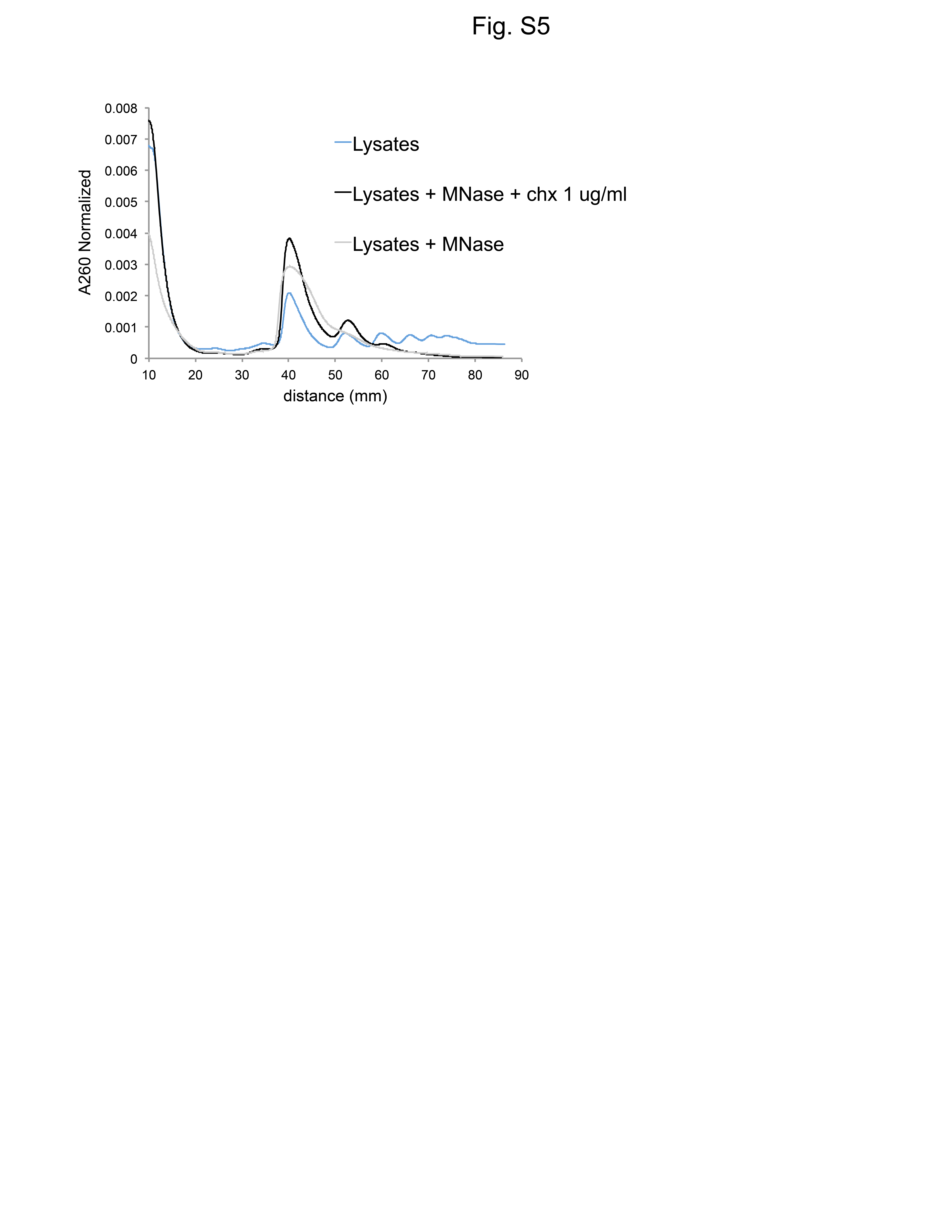

**Fig. S5:** Sucrose gradients of undigested lysates from cells with low dose cycloheximide treatment (blue), MNase digested lysates from cells with no drug treatment (gray), and MNase digested lysates from cells with low dose cycloheximide treatment (black).

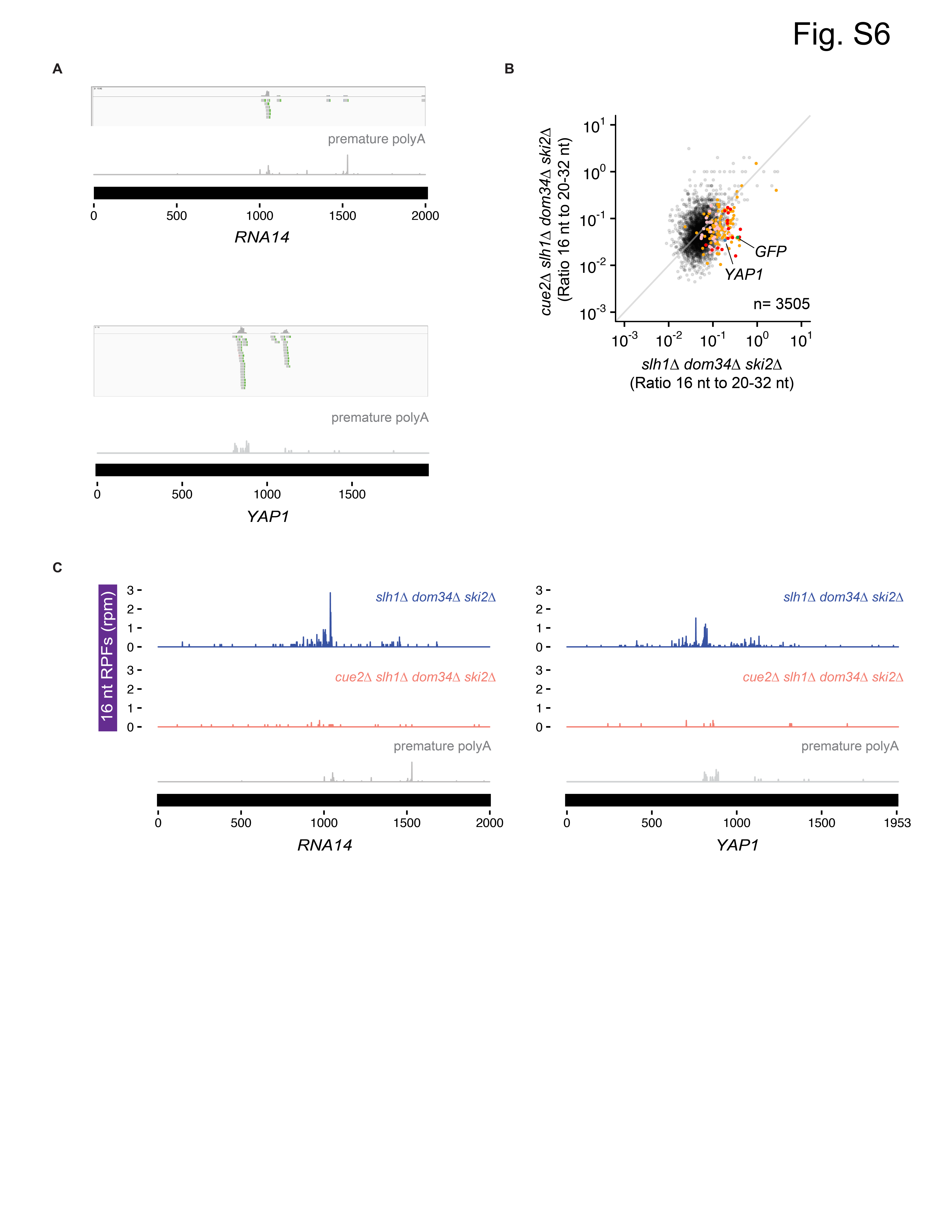

**Fig. S6:** Cue2-dependent 16 nt RPFs on prematurely polyadenylated mRNAs. **(A)** Examples of reads with untemplated A’s on *RNA14* (top) and *YAP1* (bottom). **(B)** Ratio of 16 nt over 20-32 nt RPFs plotted from *slh1∆ dom34∆ ski2∆* and *cue2∆ slh1∆ dom34∆ ski2∆* strains. Genes in orange indicate those with reproducibly decreased 16 nt RPFs upon *CUE2* deletion. Pink dots indicate genes with premature polyadenylation identified empirically. NGD-CGA reporter, *YAP1* and *RNA14* are labeled and shown in green, blue and purple, respectively **(C)** Examples of 16 nt RPFs mapped to genes, *RNA14* (left) and *YAP1* (right) in *slh1∆ dom34∆ ski2∆* and *cue2∆ slh1∆ dom34∆ ski2∆* strains with known premature polyadenylation sites (grey traces, from Pelechano et al. 2013).
